# CuBe: a geminivirus-based copper-regulated expression system suitable for post-harvest activation

**DOI:** 10.1101/2024.04.30.591823

**Authors:** Elena Garcia-Perez, Marta Vazquez-Vilar, Rosa Lozano-Duran, Diego Orzaez

## Abstract

The growing demand for sustainable platforms for biomolecule manufacturing has fueled the development of plant-based production systems. Agroinfiltration, the current industry standard, offers several advantages but faces limitations for large-scale production due to high operational costs and batch-to-batch variability. Alternatively, here, we describe the CuBe system, a novel bean yellow dwarf virus (BeYDV)-derived conditional replicative expression platform stably transformed in *Nicotiana benthamiana* and activated by copper sulfate (CuSO_4_), an inexpensive and widely used agricultural input. The CuBe system utilizes a synthetic circuit of four genetic modules integrated into the plant genome: (i) a replicative vector harboring the gene of interest (GOI) flanked by cis-acting elements for geminiviral replication and novelly arranged to enable transgene transcription exclusively upon formation of the circular replicon, (ii) copper-inducible Rep/RepA proteins essential for replicon formation, (iii) the yeast-derived CUP2-Gal4 copper-responsive transcriptional activator for Rep/RepA expression, and (iv) a copper-inducible Flp recombinase to minimize basal Rep/RepA expression. Copper sulfate application triggers the activation of the system, leading to the formation of extrachromosomal replicons, expression of the GOI, and accumulation of the desired recombinant protein. We demonstrate the functionality of the CuBe system in *N. benthamiana* plants expressing high levels of eGFP and an anti-SARS-CoV-2 antibody upon copper treatment. Notably, the system is also functional with post-harvest copper application, a strategy with potential advantages for large-scale biomanufacturing. This work presents the CuBe system as a promising alternative to agroinfiltration for cost-effective and scalable production of recombinant proteins in plants.

## INTRODUCTION

Manufacturing recombinant biomolecules in plants is becoming a flourishing new bioindustry (Eidenberger et al., 2023). In the last two decades, *Agrobacterium*-mediated transient expression in *Nicotiana benthamiana* by agroinfiltration has become the industry standard for recombinant protein production in plants (Chung et al., 2022; Shanmugaraj et al., 2020; Wani & Aftab, 2022). Product yields obtained with agroinfiltration can be further increased with upstream strategies incorporating viral replicons. These strategies involve the delivery of the gene of interest imbibed in a minimal viral unit capable of self-replicating in the plant cell. The MagnIcon (Marillonnet et al., 2005) and TRBO (Lindbo, 2007) systems are two noteworthy examples of replicative systems for agroinfiltration based on the single-stranded RNA (ssRNA) tobacco mosaic virus (TMV), both producing remarkable yields of recombinant proteins.

While RNA virus-based vectors can offer high titers for transient recombinant protein expression, they have significant limitations. Importantly, they can only integrate inserts of a limited size without disrupting RNA replication, and the absence of proofreading mechanisms in RNA-dependent RNA polymerases reduces transcript fidelity (Ahlquist et al., 2005; Castro et al., 2005; Sainsbury & Lomonossoff, 2008). Furthermore, superinfection exclusion in RNA viruses prevents using the same vector to deliver multiple transgenes (Folimonova et al., 2010; Gleba et al., 2007; Julve et al., 2013).

In contrast to ssRNA viruses, single-stranded circular DNA viruses like geminiviruses handle larger insertions than RNA viral vectors and do not present homologous (or replicative) interference for the co-expression of multiple proteins (Chen et al., 2011). Mor et al. (2003) pioneered the generation of a geminiviral vector for agro-delivery based on bean yellow dwarf virus (BeYDV) (Liu et al., 1998). The minimum viral elements required for BeYDV replication are the long and the short intergenic regions (LIR and SIR, respectively) and the Replication-associated protein (Rep) and RepA. Geminiviral replicons employed in agroinfiltration basically comprise a gene of interest (GOI) flanked by an upstream LIR and a downstream element comprising a SIR and a LIR sequence (LSL vector) (Zhang & Mason, 2006). The Rep/RepA bicistronic gene involved in replicating the geminiviral vector can be co-delivered to the plant in a different binary vector or embedded in the replicative vector between SIR and LIR (Chen et al., 2011). Vaccines, antibodies, enzymes, and virus-like particles have been expressed through BeYDV and other geminivirus-based systems (Bhattacharjee & Hallan, 2022; Chen et al., 2011). Recently, the BeYDV technology produced an antibody in *N. benthamiana* that was successfully tested *in vivo*, showing potential as an immunotherapeutic drug (Rattanapisit et al., 2023).

Despite its many advantages, agroinfiltration also entails severe limitations for large-scale bioproduction, mainly concerning production costs associated with scaling up (Fischer et al., 2012; Schillberg et al., 2019). Some studies estimate that agroinfiltration represents up to 7.5 % of the operational costs (Buyel & Fischer, 2012), plus the additional capital costs required for bacterial growth, including fermenters, infiltration tanks, and segregated infiltration rooms. Also, transient expression implies a high degree of heterogeneity in the process of vector delivery, leading to batch-to-batch variation that hinders compliance with Good Manufacturing Practices (GMP) (Gleba et al., 2013; Schillberg et al., 2019). Furthermore, the presence of *Agrobacterium* in the resulting biomass makes downstream processing difficult for specific sensitive applications. Therefore, alternative systems are required to take full advantage of the safety and scalability inherent to agricultural platforms as required by the increasing demand.

One of the possible directions to overcome agroinfiltration shortcomings is the engineering of genome-integrated, conditionally activated replicons. These are silent virus-derived replicative units integrated into the plant genome, which become active for replication only upon activation by a specific trigger. Conditional replicative platforms resemble transient expression systems in that recombinant products only accumulate at the production phase without interfering with plant development and minimizing gene silencing from long-term transgene overexpression (Mortimer et al., 2015). Additionally, an inducible system allows the expression of cytotoxic proteins in the plant without penalty in plant growth (Dugdale et al., 2013). Unfortunately, few examples exist for stable replicons due to the difficulties of engineering conditional systems that efficiently maintain the replicon in a silent state in the absence of the trigger. Werner et al. (2011) developed an ethanol-inducible system based on the MagnIcon TMV strategy. To achieve tight control of replication, the viral vector was deconstructed in a replicon module and a cell-to-cell movement module, with both modules being placed under the control of an inducible promoter.

Compared to ssRNA-based systems, the stable integration of geminivirus-based replicons has proved more amenable. Zhang & Mason (2006) stably transformed potato plants with a BeYDV-based LSL vector in which the BeYDV Rep gene was regulated by an alcohol-inducible promoter (Caddick et al., 1998). In this system, the BeYDV pro-replicon carried a fully functional transcriptional unit (TU) carrying the GOI (GFP), leading to high levels of basal expression. In 2013, Dugdale et al. developed transgenic tobacco plants harboring the In Plant Activation (INPACT) system based on the tobacco yellow dwarf virus (TYDV). In the INPACT pro-replicon design, the GOI was split into two pieces and only reconstituted upon circularization through the action of ethanol-induced TYDV Rep/RepA proteins. To our knowledge, all plant-made conditional replicons engineered for molecular farming processes to this date have employed ethanol as the trigger molecule. Whereas ethanol has low toxicity compared with other triggers employed in plant research, such as glucocorticoids, its use in large-scale installations is problematic due to its relatively high cost, high flammability, volatility, and the requirements of special permissions for its handling (Ma et al., 2021).

In this work, we describe the development of the CuBe system, a conditional BeYDV-derived replicative expression system activated by the Cu^++^ ions of copper sulfate (CuSO_4_), an inexpensive chemical trigger widely employed in agriculture, including organic agriculture, for crop protection (Lamichhane et al., 2018; Tamm et al., 2022). The new system is based on the mild strain of BeYDV (Regnard et al., 2010) and the yeast-based copper responsive factor CUP2-Gal4, which acts on a Copper Binding Site (CBS) DNA operator (Garcia-Perez et al., 2022). Four genetic modules are combined in the plant genome to get a fine-tuned geminiviral control of recombinant protein expression: the LSL replicative vector, the copper-inducible Rep/RepA proteins, the copper-dependent transcriptional activator CUP2-Gal4, and the copper-inducible Flp recombinase. The LSL replicative vector, here called Geminino, harbors a transcriptional unit (TU) of interest where the promoter is downstream of the GOI and transcriptional terminator. Due to this configuration, the GOI is only expressed from extrachromosomal circular replicons released from the plant chromosome by the Rep protein: upon release, the promoter relocalizes upstream of the GOI in the circular replicative configuration, and the effective transcription can proceed. The expression of the bicistronic Rep/RepA is placed under the transcriptional regulation of the copper-responsive factor CUP2-Gal4. To minimize the basal expression levels of Rep/RepA, a transcriptional terminator flanked by the FRT recombination sites (Lloyd et al., 2022) was embedded into the copper-inducible promoter of Rep/RepA. Upon copper-induced Flp recombinase expression, the terminator flanked by the FRT sites is removed, and Rep expression can occur. The combination of all four modules forms a synthetic gene circuit that enables refined plant and viral engineering upon induction by an agronomically friendly inducer like copper. Here, we show the development of two *N. benthamiana* plant lines conditionally expressing high levels of eGFP and an anti-SARS-CoV2 antibody following different copper application regimes. Interestingly, we also show that the system is suitable for post-harvest activation regimes, a strategy with interesting implications for large-scale manufacturing.

## RESULTS

### Optimization of the replicative module for the CuBe system

The functioning model for a BeYDV-derived LSL vector (pro-replicon) is depicted in Figure 1a. In the search for an optimal design of the CuBe system with minimal basal expression, the standard sequential order of the transcriptional unit within the pro-replicon (hereafter referred to as standard configuration) was altered, creating alternative configurations where the transcription of the GOI is impeded in the lineal pro-replicon form, but enabled upon circularization. This strategy was earlier proposed by Dugdale et al. (2013) for the so-called INPACT system, where the splitting of the transcriptional unit takes place at the CDS level, leaving a spliceable intron in between, containing the LIR (Figure 1b). Here, we modified the original INPACT configuration to facilitate golden gate-based modular cloning methods (Engler et al., 2014). In the so-called Geminino 1.0 configuration, a leader intron was placed in the 5’UTR, moving the splitting site and its associated cloning scar outside the CDS, and thus facilitating the cloning of standard DNA elements (Sarrion-Perdigones et al., 2013). In the Geminino 2.0 design, the intron was located immediately after the start codon (ATG), also avoiding the truncation of the CDS, but reducing chances of spurious CDS expression in the pro-replicon linear state.

**Figure 1.**
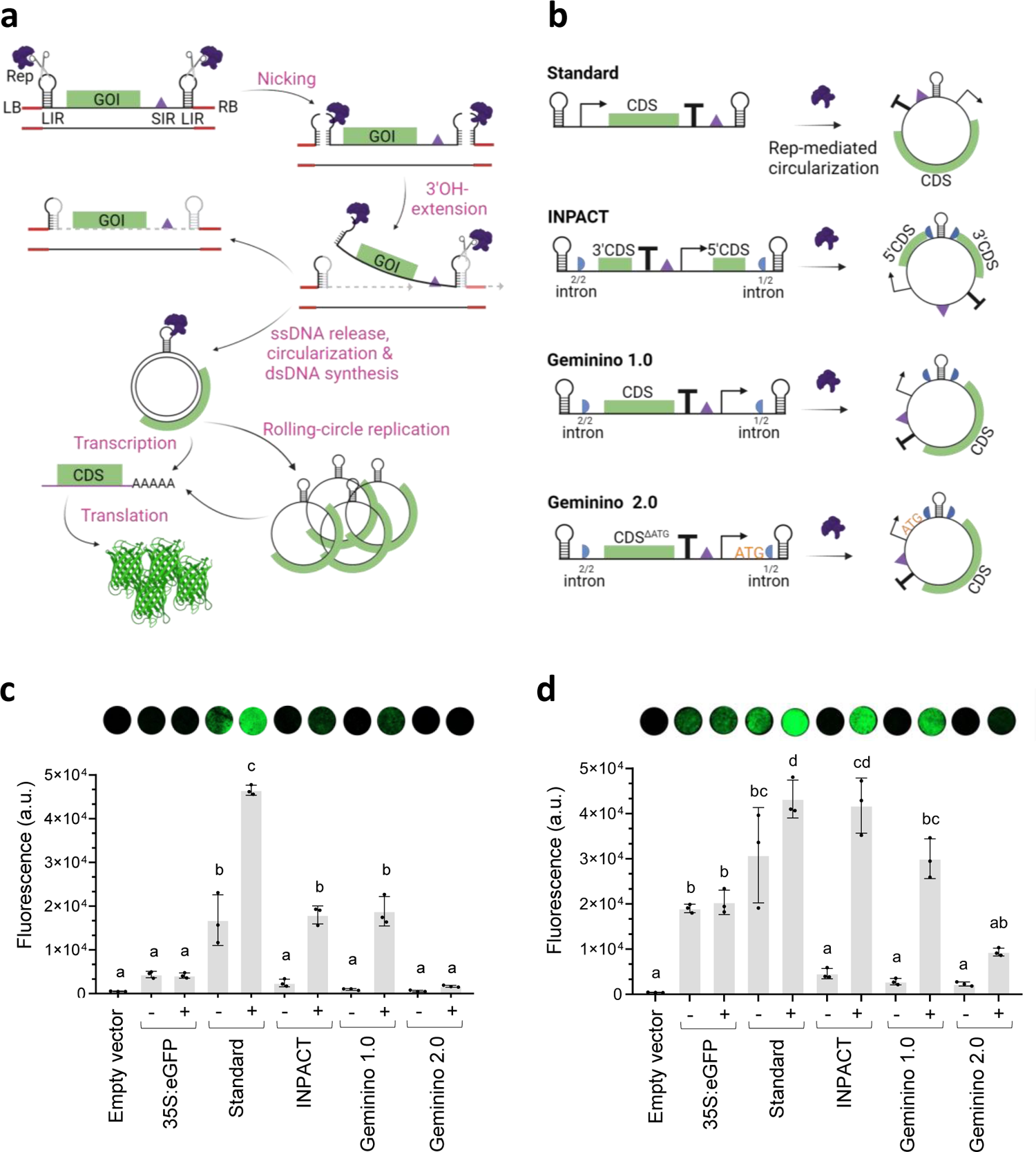
Design and evaluation of the CuBe vector for regulated protein expression. **a)** Schematic representation of the functioning model of the BeYDV-derived replicative system. The gene of interest (GOI) is flanked by long and short intergenic regions (LIR and SIR, respectively) configuring the LSL vector (pro-replicon) that is embedded between the left and right borders (LB and RB, respectively) of the T-DNA. The BeYDV Replication-associated protein (Rep) nicks the conserved region in LIR and triggers the vector’s circularization, enabling plant cell machinery to transcribe and replicate it. **b)** Schematic representation of CuBe vector configurations. The coding sequence (CDS) of interest is rearranged with the viral elements to control pre-circularization expression levels. **c)** Transient expression of enhanced Green Fluorescent Protein (eGFP) driven by different CuBe vector configurations. All constructs were introduced into *Nicotiana benthamiana* leaves via agroinfiltration. Empty vector is the negative control, 35S:eGFP is the non-replicative control, and standard, INPACT, Geminino 1.0, and Geminino 2.0 represent configurations in b). The minus (-) and plus (+) signs indicate the absence or presence of Rep/RepA co-expressed with eGFP, respectively. **d)** Same experiment as in **c)** but including the silencing suppressor p19 from *Tomato bushy stunt virus* (TBSV). Error bars represent standard deviation (n=3). Statistical analysis was performed using one-way ANOVA with Tukey’s multiple comparisons test (P-value ≤ 0.05). Variables with the same letters belong to the same statistical group. Figure includes images from Biorender (biorender.com).

The enhanced Green Fluorescence Protein (eGFP) gene was used as the reporter to assay the functionality of each configuration. All four BeYDV-based vectors were agroinfiltrated in *N. benthamiana* leaves in the presence or absence of Rep/RepA, and fluorescence was measured three days post infiltration (dpi). As seen in Figure 1c, the standard configuration produced the strongest signal, although with high background levels in the absence of Rep/RepA. In contrast, all three circularization-dependent configurations showed low fluorescence levels in the absence of Rep/RepA. INPACT and Geminino 1.0 configurations were strongly activated upon Rep/RepA-mediated circularization, whereas Geminino 2.0 levels remained at basal levels. The co-expression of the silencing suppressor p19 raised fluorescence levels in most configurations, including Geminino 2.0, which was then detectable above background levels (Figure 1d). Based on these results, we decided to select Geminino 1.0 for developing a modular and tightly controllable replicative expression system based on BeYDV.

### Optimization of a copper-regulated sensor module for the CuBe system

Owing to its agronomically-friendly characteristics, we selected CuSO_4_ as the chemical trigger for Rep/RepA activation and subsequent pro-replicon circularization. For this purpose, we adapted a plant copper sensor previously optimized in our laboratory (Garcia-Perez et al., 2022). The initial sensor, based on single copper regulation, comprised two transcriptional units: the first TU (pNOS:CUP2-Gal4) harbored the copper-dependent transcriptional factor CUP2-Gal4 driven by a constitutive pNOS promoter; in the second TU (CBS:minDFR:Rep/RepA), the CBS operator linked to the minDFR minimal promoter was coupled to the BeYDV Rep/RepA CDS. We first tested the ability of the copper sensor to activate a Geminino 1.0-eGFP pro-replicon in transient assays (depicted in Figure 2a). As can be observed in the red dashed line box in Figure 2b, this single-regulated circuitry resulted in copper activation of eGFP with relatively low background levels. Next, constructs containing the sensor module (comprising CBS:minDFR:Rep/RepA and pNOS:CUP2-Gal4 TUs) and the pro-replicon module (Geminino 1.0-eGFP) were stably co-transformed into *N. benthamiana* plants. However, despite our attempts, this strategy was unsuccessful. Only six regenerated shoots were obtained from 120 transformed *N. benthamiana* leaf discs, one of which lacked the CUP2-Gal4 gene, while another one was negative for Rep/RepA. From the remaining four fully equipped transgenic *N. benthamiana* plants, only one showed GFP expression but failed to produce viable seeds and proved unrecoverable by *in vitro* regeneration (Table S1). A parallel experiment with tobacco plants showed similar setbacks (data not shown).

**Figure 2.**
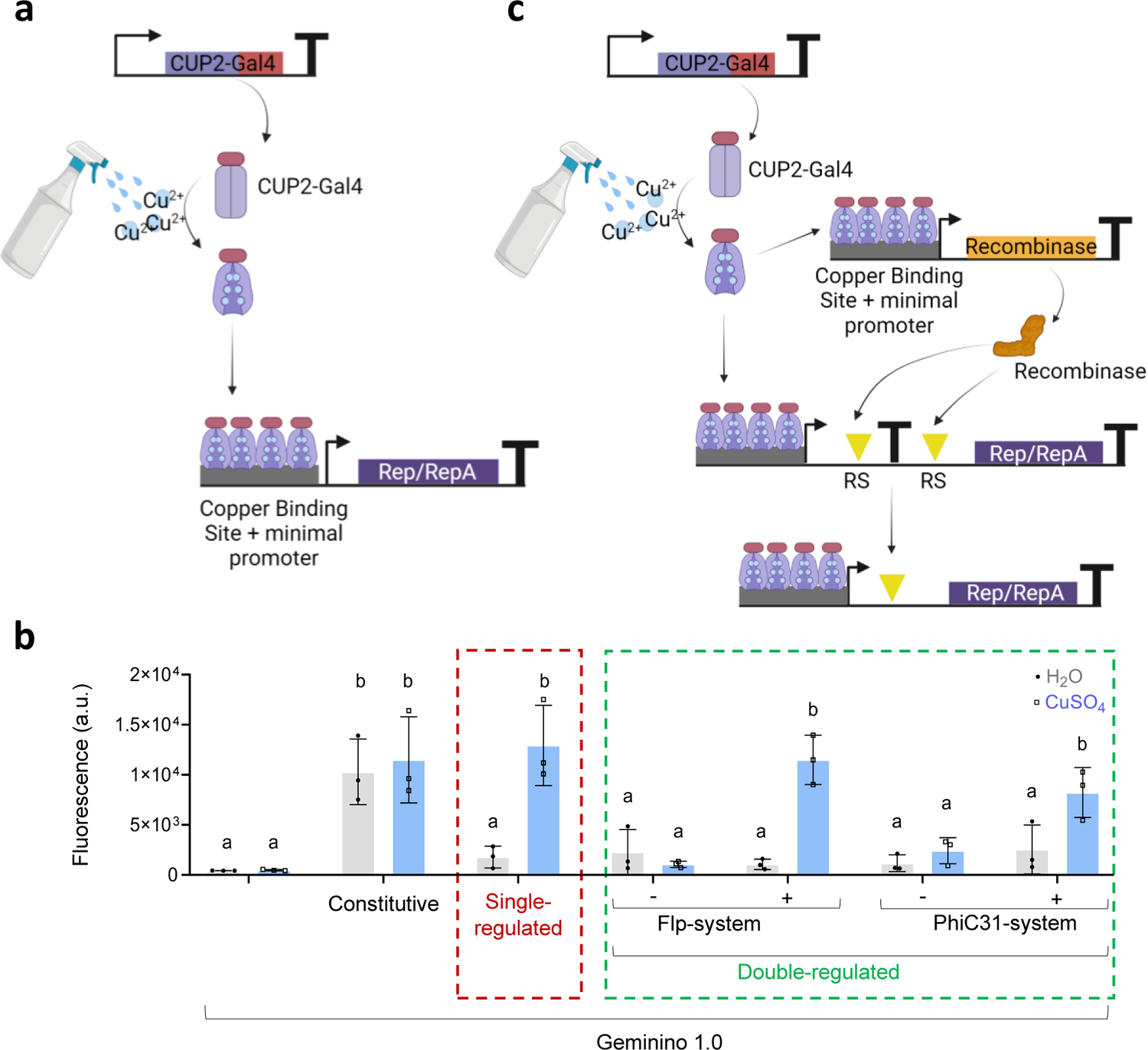
Design and evaluation of copper-mediated regulation of BeYDV Rep/RepA proteins in the CuBe system. **a)** Schematic representation of the single copper-inducible expression of BeYDV Rep/RepA proteins. The CUP2 fused to the activation domain Gal4 changes its conformation upon copper binding and binds a Copper Binding Site (CBS) operator, allowing transcription of the downstream Rep/RepA gene. **b)** Transient expression levels of enhanced Green Fluorescent Protein (eGFP) in the Geminino 1.0 vector using different regulatory systems: pNOS-driven Rep/RepA expression (Constitutive), single-layer copper regulation as depicted in a) (Single regulated); double-layer of copper regulation as is depicted in c) (Double-regulation). Both the Flippase (Flp) and the PhiC31 systems were assayed in the presence (+) or absence (-) of the respective recombinase construct. All constructs were introduced into *Nicotiana benthamiana* leaves via agroinfiltration. Fluorescence was measured in leaf discs in water (H_2_O) or 0.1 mM copper sulfate (CuSO_4_). **c)** Schematic representation of the double-regulated system. A copper-regulated recombinase acts on the Recombination Sites (RS), flanking a transcriptional terminator (represented as a T), allowing for the de-repression of Rep/RepA transcription. Error bars represent standard deviation (n=3). Statistical analysis was performed using one-way ANOVA with Tukey’s multiple comparisons test (P-value ≤ 0.05). Variables with the same letters belong to the same statistical group. Figure includes images from Biorender (biorender.com).

The limitations observed in developing transgenic plants carrying copper-inducible geminiviral elements led us to speculate that basal levels of Rep/RepA may interfere with plant development. To address this concern, we introduced, as an additional layer for Rep/RepA regulation, a memory switch mediated by site-specific recombination, as earlier proposed by Lloyd et al. (2022) (Figure 2c). An OCS transcriptional terminator (OCSt) flanked by Recombination Sites (RS) was inserted in the minDFR, upstream of the Rep/RepA coding sequence. Following this general configuration, two recombination systems were assayed: the Flp recombinase in combination with its respective FRT recognition targets, and the PhiC31 recombinase combined with its attB and attP target sites. In both systems, the expression of the recombinases was also mediated by the CBS:minDFR promoter (Figure 2c), aiming at reducing replicon activation in the absence of copper treatment.

The performance of the new double-layer regulated sensor modules (employing CBS:FRT:OCSt:FRT:minDFR:Rep/RepA or CBS:attB:OCSt:attP:minDFR:Rep/RepA switches respectively) was compared to the previous, single-layer regulation cassette (employing the CBS:minDFR:Rep/RepA construct) in the transient experiments shown in Figure 2b. Fluorescence measurements showed that eGFP expression was low in the absence of copper for all three copper-regulated strategies. Upon copper induction, fluorescence levels in all three systems raised up to levels equivalent to those obtained when employing a constitutive Rep/RepA expression construct (pNOS:Rep/RepA). Interestingly, the Flp system showed lower background levels, and it was therefore selected for a second attempt to stably integrate the CuBe system in *N. benthamiana*.

### Stable *N. benthamiana* CuBe-eGFP plants

The optimized CuBe gene circuit was stably transformed in *N. benthamiana* following a co-transformation approach involving two T-DNA multigene constructs: the so-called sensor module, on the one hand, encoded the Rep/RepA, the Flp, and the CUP-Gal4; the processor module on the other hand (a.k.a. amplification or pro-replicon module) comprised the Geminino1.0-eGFP cassette (Figure 3a). The segregation of the system into two autonomous modules allows for the independent segregation of each module with the potential to generate new functional circuits through super-transformation or sexual crosses. To facilitate co-transformation, the two modules were cloned into two binary vectors with compatible replication origins. The amplification module was cloned into a mini binary vector, pLX-B3, with replication origin pBBR1 (Pasin et al., 2017). The sensor module was included in a pDGB3 vector (Vazquez-Vilar et al., 2017) based on pCAMBIA containing the replication origin pVS1. Both vectors were simultaneously used in a two-plasmid/one-*Agrobacterium* strain strategy for multiple delivery of T-DNAs to the plant cell. Furthermore, several additional elements were introduced in the final constructs to improve stability and facilitate the proper regulation of transgene in the genomic context. In the sensor module, each transcription unit was separated by intergenic regions derived from NbDFR homeologous genes. In the pro-replicon module, the Geminino 1.0-eGFP was flanked by two matrix attachment regions (MARs) from *Arabidopsis thaliana* chromosome 4 (MAR10) (Pérez-González & Caro, 2019). After being successfully tested in transient assays (Figure 3b), the transformation-ready constructs were co-transformed into *N. benthamiana* plants are aiming at creating first-generation CuBe plants, as graphically depicted in Figure 3c. From this co-transformation experiment, eleven T0 plants were recovered and analyzed in genetic screening, aimed at detecting the presence of the two genetic modules (Table S2), and in functional analysis, aimed at detecting eGFP in leaf discs treated with CuSO_4_ (Figure 3d). All regenerated plants contained both the sensor and processor modules. Surprisingly, careful examination of the sensor module revealed that the OCS terminator (OCSt) originally interrupting Rep/RepA transcription was lost in all the eGFP-expressing lines except for line 9 during the transformation/regeneration steps. Plants 1 and 2 (non-functional) and plant 9 (low eGFP expression) were the only ones maintaining the OCS terminator (Figure 3d). We speculate that the basal expression of Flp during the plant regeneration or gametic formation may have led to the excision of the OCS terminator. Despite the absence of one of the elements of the Flp memory switch, T0 plants showed proper copper inducibility in leaf disc assays (Figure 3d). Therefore, we proceeded with the functional assessment of the next generation.

**Figure 3.**
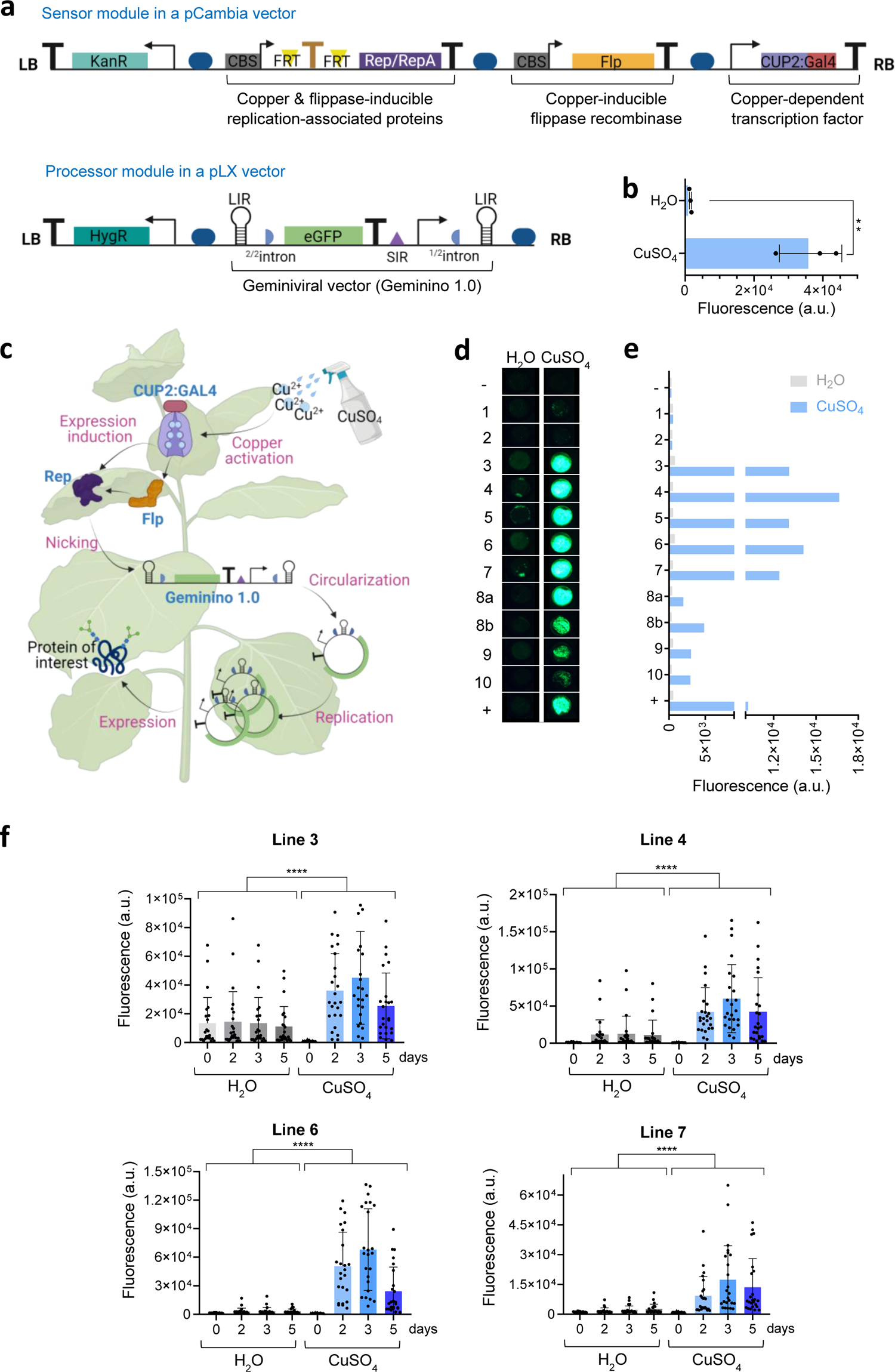
Design and evaluation of the CuBe system in transgenic *Nicotiana benthamiana*. **a)** Schematic representation of the two gene modules used for stable plant transformation. Flp is the flippase recombinase; CUP-Gal4 is the copper-dependent transcriptional factor; KanR is the gene nptII conferring kanamycin resistance; HygR is a hygromycin resistance gene. Arrows indicate transcriptional promoter regions; ‘T’ indicates transcriptional terminators; the golden ‘T’ is the OCS T flanked by flippase target sites (FRT); CBS is the copper binding site recognized by CUP2-Gal4; LIR and SIR are intergenic regions of BeYDV; the blue circular shapes are different intergenic regions acting as gene insulators; half circles are the initial (1/2 intron) and the final (2/2 intron) segments of a unique intron; LB and RB are Left and Right Borders of T-DNA, respectively. **b)** Transient evaluation of constructs in a) by agroinfiltration of *N. benthamiana* leaves and fluorescence measurement of leaf discs incubated either in water (H_2_O) or 0.1 mM copper sulfate (CuSO_4_) (n=3). **c)** Schematic representation of the functioning of the CuBe system once stably integrated into the plant. **d)** Fluorescence records at 3 days post-incubation of leaf discs from CuBe-eGFP T0 plants in H_2_O or 0.1 mM CuSO_4_. **e)** Quantification of fluorescence levels shown in d). **f)** Time-course of eGFP activation in leaf discs taken from T1 CuBe plants upon incubation in H_2_O or 0.1 mM CuSO_4_. Each point represents the fluorescence levels of an individual disc taken from a different T1 plant (n=24). Error bars represent standard deviation. Statistical analysis was performed using a T-student (P-value ≤ 0.05) for b) and a one-way ANOVA with Tukey’s multiple comparisons test (P-value ≤ 0.05) for f). For the ANOVA test in f), the areas under the curve of the H_2_O and CuSO_4_ time-course treatments were compared. Asterisks represent statistical significance. Figure includes images from Biorender (biorender.com).

Based on optimal regulatory performance (Figure 3e), we chose four T0 plants for progression to the first transgenic generation (T1 lines 3, 4, 6 and 7). In addition, the offspring of line 9 was also analyzed, resulting in a mixed population of switchable and non-switchable plants, the latter group maintaining the OCS terminator (Figure S1). Functional analysis of 24 plants from each selected T1 line is shown in Figure 3f. Induction of eGFP was observed in all four lines after two days of copper treatment. Segregation analysis of transgene modules in the four T1 lines revealed that all lines contained a single copy of the sensor module (Tables S3 and S4). However, lines 3 and 6 contained two copies of the pro-replicon module, whereas lines 4 and 7 contained a single copy. To facilitate the interpretation of results in the subsequent analysis aimed at optimizing treatment conditions, single-copy lines 4 and 7 were selected to progress to the T2 generation.

### Copper treatment regimes

Homozygous T2 CuBe-eGFP lines 4.11 and 7.8 (Table S5) were next assayed using different copper treatment strategies, aiming at establishing easily scalable induction regimes that could reduce operational costs in both contained and open-field conditions. We initially analyzed a spray application of plants with a CuSO_4_ solution combined with 0.05% fluvius surfactant. Interestingly, 72h after treatment, widespread fluorescence was observable throughout the surface of the leaves, indicating appropriate CuSO_4_ uptake. We experimentally determined optimal copper concentrations for spraying at 5 mM CuSO_4_ (Figure 4a). Maximum eGFP expression was observed after 4 days of one-time 5 mM CuSO_4_ application (Figure 4b), the time point at which we stopped the eGFP measurement due to the clear presence of necrosis in leaves, possibly due to copper toxicity (Figure S2).

**Figure 4.**
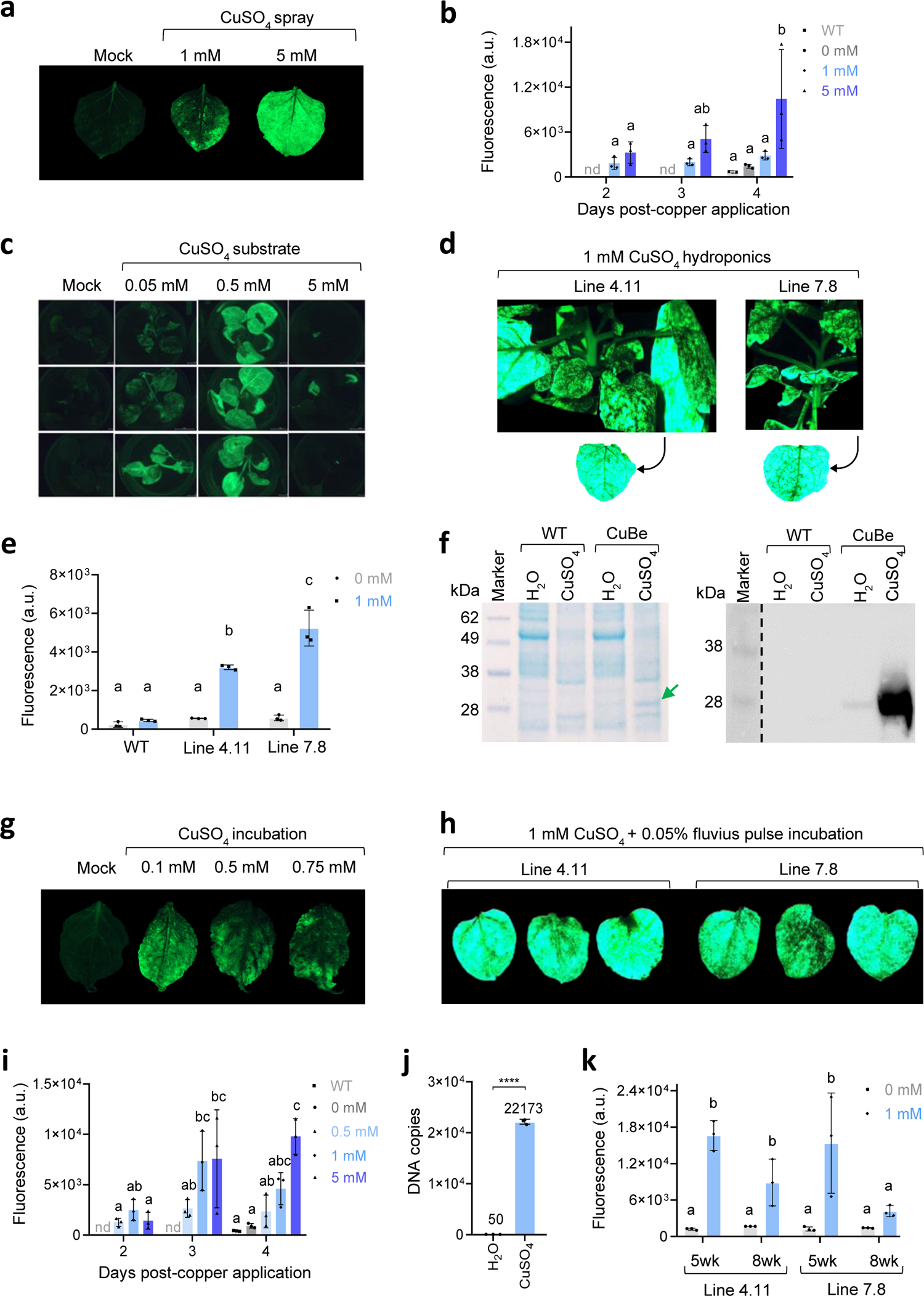
Copper induction analyses of stable *Nicotiana benthamiana* CuBe-eGFP plants. **a)** Leaves of CuBe-eGFP plants sprayed with H_2_O or copper sulfate (CuSO_4_) and visualized at the fluorescent magnifier after 3 days post application (dpa). **b)** Time course of fluorescence emitted from sprayed leaves of CuBe-eGFP plants treated with different CuSO_4_ concentrations. Each data point represents an independent leaf. **c)** Fluorescence reported at 5 dpa of CuBe-eGFP seedlings grown in agar media. **d)** Fluorescence image of plants grown in 1mM CuSO_4_ hydroponics at 3 dpa. **e)** Quantification of fluorescence emitted in plant leaves from wildtype (WT) and transgenic lines 4.11 and 7.8 treated as in d). Each data point represents an independent plant. **f)** Coomassie-stained SDS-gel electrophoresis of total soluble proteins (left panel) and Western blot of the same samples (right) from WT and 7.8 CuBe-eGFP plants induced hydroponically as in d). **g)** Representative fluorescence images of CuBe-eGFP leaves after 2 days of continuous post-harvest incubation in CuSO_4_. **h)** Representative fluorescence images of CuBe-eGFP leaves pulse-treated (5 min) after harvest with a CuSO_4_ + 0.05% fluvius solution and kept in H_2_0 for 2 days. **i)** Time-course of fluorescence levels in leaves after post-harvest pulse treatments with different CuSO_4_ concentrations. **j)** DNA copy number estimation of Geminino 1.0 vector per plant haploid genome in CuBe-eGFP plants. Each point represents a qPCR technical replicate coming from three post-harvested leaves pulse-treated with 1 mM CuSO_4_ + 0.05% fluvius and kept in H_2_0 for 3 days. (n = 3). **k)** Influence of plant development stage in the accumulation of eGFP. 5wk indicates 5-weeks-old plants and 8wk indicates 8-weeks-old plants. Nd stands for no data. Each data point represents a leaf from a different CuBe-eGFP plant. Error bars represent standard deviation (n=3). Statistical analysis was performed using a one-way ANOVA with Tukey’s multiple comparisons test (P-value ≤ 0.05). Variables with the same letters belong to the same statistical group. A T-student (P-value ≤ 0.05) was used for j). Figure includes images from Biorender (biorender.com).

Next, we aimed to investigate the possibility of employing systemic CuSO_4_ treatments for the induction of the CuBe system. We first conducted a small-scale assay using seedlings grown in MS-agar plates to monitor copper movement through the plant. Seedlings were placed in MS-agar media with 0.05, 0.5, and 5 mM CuSO_4_ (Figure S3). Remarkably, as shown in Figure 4c, a systemic copper signal was successfully established, leading to strong eGFP activation in distal leaves. Two days after copper application in agar, necrosis started in CuBe seedlings at 5 mM, and no fluorescence was reported at any recorded time point (Figure S4). Seedlings in 0.05 mM did not show necrosis nor fluorescence activation. Seedlings at 0.5 mM started to show systemic fluorescence emission on day 3 post-copper application, and they did not show necrosis symptoms except localized brown spots on day 5 (Figure S3). On day 6, after changing seedlings to new agar media without copper, the new emerging leaves did not show fluorescence, indicating that eGFP expression is totally dependent on copper presence (Figure S4).

We then attempted CuSO_4_ systemic activation by watering plants grown in pots. However, all our efforts to reproduce activation in these conditions were unsuccessful. As an alternative strategy for large-scale systemic induction, we then assayed copper treatments in plants grown in hydroponic conditions by transferring the plants from a standard hydroponic media to an equivalent media supplemented with 1 mM CuSO_4_. After three days in the CuSO_4_-enriched hydroponic media, strong and widespread fluorescence levels were recorded in plant leaves, indicating successful systemic movement of the copper signal in hydroponic conditions (Figures 4d and 4e). This systemic movement of copper through the plant also produced toxicity symptoms, causing browning of roots and leaf curling. However, no necrosis was reported in the aerial and eGFP-producing vegetative material (Figure S5). The accumulation of eGFP protein in leaves activated systemically by hydroponic induction was estimated in Coomassie-stained gels, yielding approximately 120 µg of recombinant protein per gram of fresh weight, representing a 2.6% total soluble protein (TSP) (Figure S6). Re-evaluation by Western blot of the copper-dependent induction in hydroponic conditions showed remarkably low background levels of eGFP expression in non-induced conditions (Figure 4f). Given these results, we speculate that the failure to establish systemic activation in soil-grown plants could be due to the immobilization of copper because of interactions with substrate components, although this aspect will require further investigation.

Finally, owing to the fast response of the CuBe induction system, we became interested in assaying post-harvest induction in leaves as an easily scalable option. To this end, detached leaves were incubated in water tanks containing different concentrations of CuSO_4_ ranging from 0.01 mM to 1 mM. Using this approach, eGFP activation was low. Tissue damage was visible at 2 days post-incubation on 1 mM CuSO_4_ and turned strong on day 5 (Figure S7). Lower copper concentrations ended up in irregular and relatively low eGFP induction at the assayed concentrations (Figure 4g). Next, we modified the post-harvest strategy, incubating the detached leaves in 1 mM CuSO_4_ plus 0.05% fluvius for only 5 minutes and subsequently moving them to a water container for the remaining incubation time. As can be observed in Figure 4h, a much higher fluorescence signal was reported at 2 days post-copper incubation (dpci) than in the previously assessed methods and without signs of damage (Figure S8). For further optimization of the post-harvest induction, additional activation regimes were tested. Figure 4i shows that a maximum eGFP fluorescence was obtained at 4 dpci using a pulse treatment of 5 mM CuSO_4_. However, this regime also caused initial symptoms of necrosis on day 4 (Figure S9). Therefore, we decided to set the standard treatment for post-harvest induction as a 5-min pulse incubation in 1 mM CuSO_4_ 0.05% fluvius, followed by 3 days of incubation in water. Using these activation conditions, we calculated the number of copies of CuBe-eGFP replicon generated per haploid genome. As can be seen in Figure 4j, under induced conditions, the estimated transgene dosage reached more than 2×10^4^ copies per haploid genome, whereas background levels in uninduced conditions reached up to 50 replicon copies per genome (Figure S10). As a final test, we used this standard regime to compare the GFP levels produced by leaves harvested at different stages of plant development. Figure 4k shows the fluorescence emission from leaves taken from 5-week-old plants compared to 8-week-old (flowering) plants (see Figure S11 for leaf size comparison).

### Production of antibody n72_hγHC by the CuBe system in transgenic *N. benthamiana* plants

After optimizing the CuBe system using the eGFP reporter protein, we proceeded to test the versatility of the system by producing the single-chain anti-SARS-CoV-2 antibody n72_hγHC (Diego-Martin et al., 2020). This antibody is formed by a camelid variable heavy chain (VHH) domain that targets the receptor binding domain (RBD) of the SARS-CoV-2 spike (S) protein fused to the Fc region of a human IgG1. For developing CuBe-n72_hγHC plants, we employed the same sensor module as in CuBe-eGFP, whereas the pro-replicon module was reassembled using the n72_hγHC CDS instead of eGFP (Figure 5a). Nineteen co-transformed T0 lines were regenerated, and their functionality was screened in an ELISA test using plates coated with RBD (Figure 5b). Eleven of the 19 screened plants presented regulated production of the n72_hγHC antibody. Since a secretion signal peptide was fused to the n72_hγHC, we also analyzed the anti-RBD activity of an apoplast wash obtained from T0 plants. As shown in Figure 5c, the T0 CuBe-n72_hγHC plants secrete enough n72_hγHC to detect RBD without further processing of the plant material. Six T0 plant lines were brought to T1 generation, germinated in selective media, and re-analyzed in the ELISA test shown in Figure 5d. Remarkably, all T1 plants analyzed were positive for copper-induced production of n72_hγHC. Mendelian segregation analysis revealed that line 8 contained two copies of the Geminino 1.0 module, which may explain its higher reactivity levels towards RBD (Tables S6 and S7). Protein extracts of the T1 8.6 plant were analyzed by Western blot to confirm the tight copper-regulated expression of the n72_hγHC (Figure 5e). The Western blot analysis also indicated that leaves at different developmental stages, both fully expanded (FE) and non-fully expanded (NE), produce the antibody at similar levels. This was later confirmed by ELISA using plants treated with CuSO_4_ following a hydroponic regime, as shown in Figure 5f. Antibody tittering curves of crude and apoplast extracts reached end tittering points up to 3 x 10^-4^ (Figure 5g). In these induction conditions, the calculated average number of CuBe-n72_hγHC replicons raised from 4 to almost 1×10^4^ copies per haploid genome (Fig 5h). As shown in Figure 5i, Coomassie-stained SDS-PAGE from apoplast fluid showed a differential band attributed to n72_hγHC. Single-step protein affinity purification of the n72 hγHC from clarified protein extract produced an estimated yield of 4.5 µg/g FW purified single-chain antibody (Figure 5j and S12).

**Figure 5.**
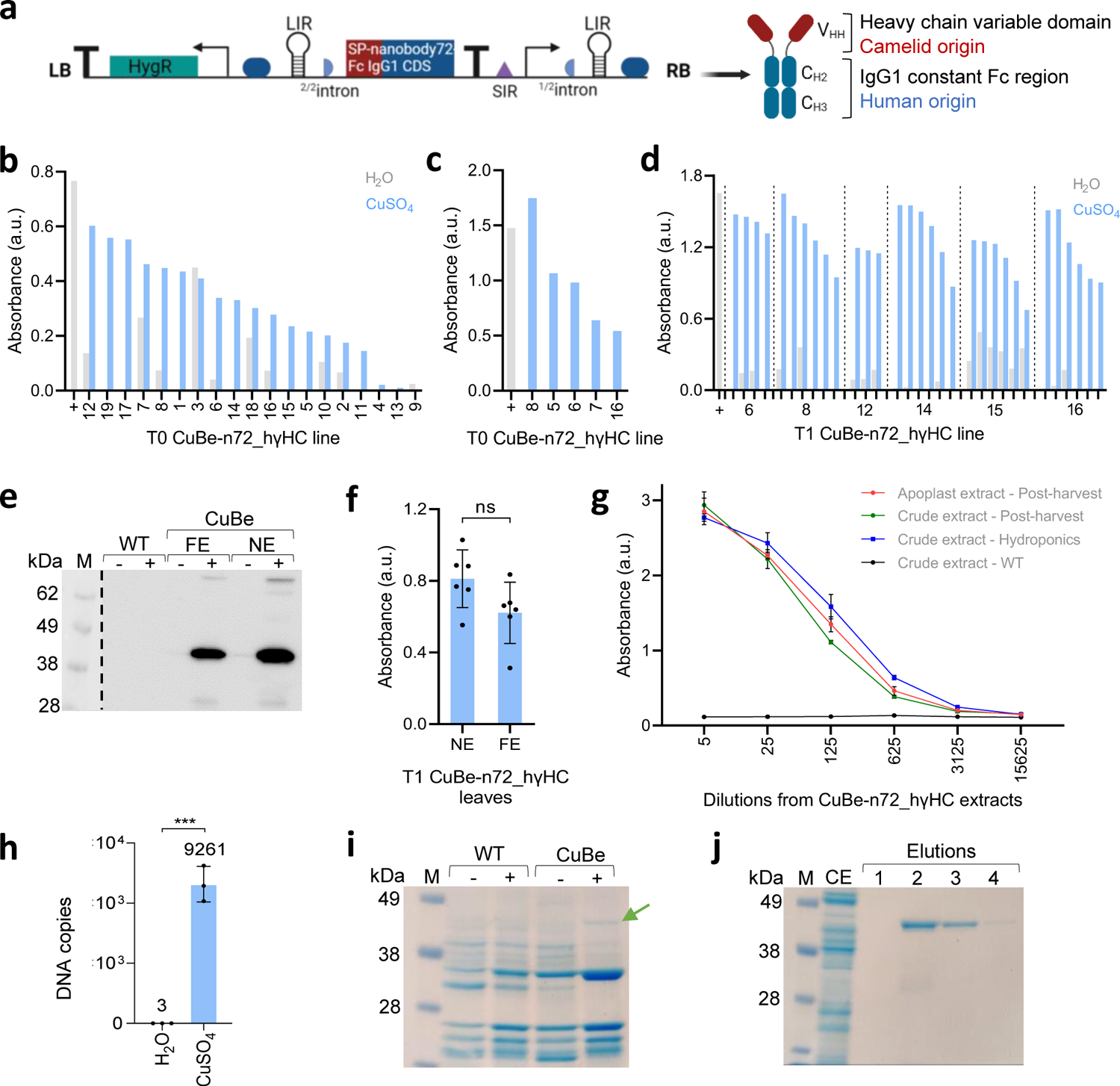
Copper-induced production of n72_hγHC antibody in stable *Nicotiana benthamiana* CuBe plants. **a)** Schematic representation of the pro-replicon module used for n72_hγHC expression. SP stands for signal peptide. HygR indicates the hygromycin resistance gene used as a selection marker. Arrows indicate promoter regions; LIR and SIR are intergenic regions of BeYDV; the blue circular shapes are intergenic regions acting as gene insulators; half circles are the initial (1/2 intron) and the final (2/2 intron) segments of a unique intron; LB and RB are Left and Right Borders of T-DNA, respectively. **b)** Antigen (RBD) ELISA screening of the T0 CuBe-n72_hγHC *N. benthamiana* plants. ELISA plates were coated with RBD and incubated with total soluble protein extracts from leaf discs of plants 1 to 19. Discs were previously incubated for 4 days in H_2_O or in 0.1 mM CuSO_4_. **c)** Antigen (RBD) ELISA of undiluted apoplast fluid obtained from selected T0 leaves incubated for 2 days in 0.1 mM CuSO_4_. **d)** Antigen (RBD) ELISA screening of different T1 CuBe-n72_hγHC plants induced with a 1 mM CuSO_4_ + 0.05% fluvius post-harvest treatment for 5 min. **e)** Western Blot analysis of total soluble protein extract from fully expanded (FE) and non-fully expanded (NF) leaves of T1 8.6 CuBe-n72_hγHC plant. Plants were treated with CuSO_4_ as indicated in d) (+) or left untreated (-). **f)** Antigen ELISA of crude extracts from different T1 plants derived from line 8, activated following a hydroponics regime with 1 mM CuSO_4_. NE indicates non-fully expanded leaves; F, full-expanded leaves. Each data point represents a leaf from a different plant. Bars represent mean ± SD, n = 6, and asterisks result from a T-student comparison (P-value ≤ 0.05). In all the ELISA bars on b), c), d), and f), the background absorbance from wild-type crude extract samples was subtracted. As a positive control (+), extract from a wildtype plant agroinfiltrated with a constitutive n72_hγHC construct. **g)** Antigen ELISA curves of apoplast and crude extracts from T1 line 8 n72_hγHC plants to compare antibody binding activities of n72_hγHC induced and extracted by different methods. The X-axis indicates the 1:5 dilution factor of the extracts from 1:5 to 1:15625. **h)** DNA copy number estimation of Geminino 1.0 vector per plat haploid genome in CuBe-n72_hγHC plants. Each point represents a qPCR technical replicate coming from three plant leaves post-harvested incubated in H_2_O or 1 mM CuSO_4_ for three days (n = 3). **i)** Coomassie-stained SDS-PAGE gel of apoplastic fluid obtained from leaves of wildtype (WT) or T1 8.6 CuBe-n72_hγHC treated with H_2_O (-) or with CuSO_4_ as indicated in d). The arrow points to the n72_hγHC antibody band size. **j)** Coomassie-stained SDS-PAGE gel showing affinity purification of n72_hγHC. M stands for molecular marker (1 kDa); CE is a crude protein extract; elutions 1-4 correspond to fractions of a protein-A purification procedure. The figure includes images from Biorender (biorender.com).

## DISCUSSION

Plants harboring stably integrated replicons offer a solution for the sustainable and cost-effective manufacture of recombinant products. Production yields can be optimized by segregating plant development from the activation of the pro-replicon. For this separation, conditional activation is required to control the timing, location, and amount of transgene expression. To date, two main transgenic systems, ‘MagnICON’ and ‘INPACT,’ excel at minimizing background expression in the absence of the inducer while maximizing protein production upon activation in plants. The TMV-based ‘MagnICON’ system (Werner et al., 2011) was implemented in *N. benthamiana* to yield up to 4.3 g GFP/kg fresh weight upon induction by a combination of root drenching and vapor of ethanol for five to seven days. The TYDV-based ‘INPACT’ system (Dugdale et al., 2013) reported yields of 25 mg GUS protein/g TSP (2.5% TSP) in *N. tabacum* or 0.1 g vitronectin/kg fresh weight in *N. benthamiana*. Our data show that yields obtained with the CuBe system, which reached up to 0.12 g GFP/kg fresh weight (2.6% TSP), are comparable to the INPACT system. The yield reported for the n72_hγHC antibody is more modest. However, antibody yields are highly dependent on variable chain sequence (Mason et al., 2012), and although not suitable for yield comparison, the CuBe-n72_hγHC plants serve as an example of the consistency, versatility, and remarkable inducibility of the CuBe system. The average 50 and 4 copies of replicon per haploid genome observed under non-induced conditions in CuBe-eGFP and CuBe-n72_hγHC lines, respectively, can be probably attributed to the occasional escape of a replicon in isolated cells (Figure S13). The accumulated evidence seems to indicate that, despite the massive increase in transgene copy number associated with the replication of geminivirus-based systems, transcription and translation levels are not as high as those of, e.g., TMV-based systems. This could be due to a lack of intrinsic gene silencing suppressor activity and/or to the absence of cell-to-cell movement that, in other viral systems, such as MagnICON, expands the expression foci in case the activation signal does not reach all the cells. Silencing in CuBe might be mitigated by silencing suppressors, as reported by Zhang & Mason (2006). On the other hand, RNA virus-based systems suffer superinfection exclusion (Folimonova et al., 2010; Gleba et al., 2007; Julve et al., 2013), a property that makes it unsuitable for producing protein complexes. Therefore, the selection between geminivirus and TMV-based systems for recombinant protein production should be guided by specific application requirements.

MagnICON and INPACT, as well as other plant conditional expression systems, rely on the ethanol-responsive AlcA:AlcR gene switch for activation (Caddick et al., 1998). However, using ethanol as an inducer entails several practical, economic, and storage drawbacks. Its high volatility makes field application practically unfeasible in open environments. Additionally, being a highly flammable compound, ethanol requires special permits for acquisition and storage tanks with high resistance to ethanol’s corrosive attack (Ma et al., 2021). One of the main advantages of the CuBe system lies in its activation by Cu^2+^ ions, which can be supplied, e.g., in the form of CuSO_4_, a low-cost agrochemical suitable for indoors and open-field applications without evaporation risks and with eco-friendly validation (Lamichhane et al., 2018; Tamm et al., 2022). Although plants use Cu^2+^ at physiological concentrations as a cofactor for a wide range of cell processes (Tsang et al., 2021), the CuBe system can be maintained in an OFF state when plants are grown on regular plant growth media, becoming activated only when intracellular concentrations reach non-physiological conditions. The origins of the Cu^2+^ sensor employed here, which derives from the yeast CUP2 system that evolved to respond only to toxic Cu^2+^ concentrations (Buchman et al., 1989), explain this behavior. Furthermore, as shown in this work, under hydroponic conditions, the plant root is not able to block the uptake of the excess copper efficiently. Consequently, the high Cu^2+^ signaling queue required for CuBe activation can be expanded systemically, allowing its transmission to distal tissues. Indeed, hydroponics stands out as an efficient method for copper conditional activation in our hands, leading to one of the highest production levels among the different application methods assayed. However, a major drawback of this method is the severe necrosis observed in the roots. It is described that exposure to high copper concentrations can lead to acute toxicity, with reduced mitotic activity in the root tips, ROS production, and nutrient uptake disruption (Küpper & Andresen, 2016). Further industrial optimizations could lead to an optimal balance between toxicity and production levels using hydroponics.

Next to copper conditional expression, CuBe incorporates other specific design features that deserve consideration in the context of inducible plant biofactories. The circuity was deliberately designed to create a triple regulatory layer comprising transcriptional activation, pro-vector circularization, and site-specific recombination. Adaptation to standardized modular cloning (Vazquez-Vilar et al., 2017) facilitated parts assembly and simplified exchange of GOIs and will eventually facilitate further vector optimization by incorporation of new regulatory DNA elements, as shown by Dusek et al. (2020). The CuBe design incorporated intergenic spacers separating different TUs to enhance the isolation of the genes of interest. We also flanked the pro-replicon by MAR regions, which are described to minimize interline variation and protect transgenes from deletions (Pérez-González & Caro, 2019). Another distinctive feature is the use of the two-plasmids/one *Agrobacterium* co-transformation strategy. The use of two separated binary vector backbones with compatible origins (Pasin et al., 2017) harboring the sensor and the amplification modules, respectively, led not only to a high transformation efficiency in our hands but, most importantly, also to independent integration of the two functional modules. In this way, once a pair of integration loci has proved optimal, offspring carrying each of the modules can be sexually segregated, and the resulting elite single-module plants can be re-used in new combinations made by super-transformation or sexual crossing.

The incorporation of a recombinase-based terminator switch (Lloyd et al., 2022), which was initially aimed at minimizing leaky pro-vector activation, turned out to facilitate the regeneration of transgenic plants harboring the CuBe system. We obtained 11 double-transformed, functional n72_hγHC transgenic plants from 90 leaf discs. We were unable to regenerate transgenic plants in previous attempts with construct designs lacking this repressor switch. Furthermore, although not fully comparable, this seems to be a higher transformation rate than that reported for sequential super-transformation in similar approaches (Dugdale et al., 2013). We favor the explanation that the terminator switch worked as a robust break, repressing leaky Rep/RepA expression during early organogenesis steps, which could otherwise lead to cell death and meristem abortion. It should be noticed that, whereas transcriptional activation follows a continuous behavior, recombinase-based regulation follows a Boolean dynamic, with only two possible statuses (active or inactive). Consequently, although terminator removal in the absence of copper seems to have happened promiscuously during plant regeneration, some cell lineages may have kept the recombinase brake sufficiently long to survive the undifferentiated stage in which other transcriptional control mechanisms are inefficient. A similar phenomenon was reported by Guiziou et al. (2023), which led to the suggestion that integrase expression could occur during gametogenesis and then be transmitted to all cells of the next generation. The loss of the terminator at early regeneration stages did not translate into copper deregulation in differentiated tissues. Interestingly, when transgenic plants maintaining the terminator switch were agroinfiltrated with a constitutive flippase, efficient removal of the terminator was observed by PCR analysis (data not shown), indicating the proper functioning of the switch. However, the same plants were not activated under copper treatment. A possible explanation for this behavior is the posttranslational gene silencing of the flippase transgene, as earlier reported for tyrosine integrases such as Flp operating in transgenic plants (Guiziou et al., 2023; R. Liu et al., 2021).

Another remarkable feature of CuBe, as compared with other commonly used conditional systems in plant biology, is the ability to launch a sustainable activation with just a pulse (five-minute) incubation in post-harvest conditions. This effect can be related to the long stability and persistence of the Cu metal ions in the plant tissues, in contrast with other highly volatile and/or more prone to degradation (organic) signaling molecules. This method can greatly simplify and reduce the cost of industrial applications, as the same CuSO_4_ solution can be used to induce multiple plant batches.

In summary, the CuBe system represents a successful integration of a potent inducible viral system into transgenic plants. It stands out from previous systems due to its copper-inducible activation, which allows scaling up plant-based protein production in open field conditions. We have demonstrated that the system activation is highly versatile and efficient both in vivo by spraying or hydroponic irrigation and ex vivo by pulse-incubation of leaves. This system has the potential to be integrated into several plant species since BeYDV can infect multiple dicotyledonous plants. The system’s flexibility in terms of activation and the repertoire of plants it could be integrated into highlight its potential as a powerful tool in the field of molecular farming.

### EXPERIMENTAL PROCEDURES

#### Cloning and assembly of the GoldenBraid constructs

All the genetic constructs used in this work were assembled using GoldenBraid (GB) (Sarrion-Perdigones et al., 2013; Vazquez-Vilar et al., 2017). All the employed expression vectors are detailed in Table S8, along with their identification labels on the GoldenBraid repository.

#### Agroinfiltration experiments

Genetic constructs were transiently expressed in wild-type *Nicotiana benthamiana*. *Agrobacterium tumefaciens* C58 harboring expression vectors were grown in LB with 50 μg/ml kanamycin/spectinomycin and 50 μg/ml rifampicin. After 16 h at 28°C/250 rpm, bacteria cultures were resuspended in infiltration buffer (10 mM MES, 10 mM MgCl2, and 200 μM acetosyringone, pH 5.6) and infiltrated at OD_600_ 0.1 into three expanded leaves of five-week-old *N. benthamiana* plants for co-expression of GB elements. Plants were maintained in a controlled environment (24°C light/20°C dark, 16h light/8h dark).

#### Fluorescence quantification of eGFP expression

Leaf discs (5 mm diameter) were collected and placed upside-down in a black 96-well plate (Corning, #3792) with 250 µl distilled water per well. Fluorescence (excitation 490 nm, emission 510-570 nm) was measured using a GloMax®-Multi Detection System (Promega). For time courses, plates were incubated at 25°C, 16h light/8h dark, and 60-70% humidity. Alternatively, 50 µl of total soluble protein extract was loaded per well and measured similarly.

#### Fluorescence imaging of eGFP expression

The fluorescence imaging of plant material was done by using a magnifier Leica MZ 16 F. The conditions for the imaging were 2-sec exposition, 10x gain, 0 saturation, and 1.04 gamma.

### Stable transformation of *Nicotiana benthamiana*

Sensor and replicative modules were co-electroporated into *A. tumefaciens* LBA4404. Saturated cultures (OD_600_ 0.2) were inoculated overnight in TY medium with antibiotics (kanamycin and spectinomycin) and 200 μM acetosyringone. Leaf discs from six-week-old surface-sterilized *N. benthamiana* leaves were placed onto co-cultivation medium (MS medium at pH 5.7, 1 mg/L 6-benzylaminopurine (BAP), 0.1 mg/L naphthalene acetic acid (NAA) and 9 g/L phytoagar) for 24 h. Explants were immersed in the *A. tumefaciens* culture for 15 minutes before being placed back onto the co-cultivation medium. After 2 days, discs were transferred to selection medium (including 100 mg/l kanamycin, 20 mg/l hygromycin, 200 mg/l carbenicillin) for shoot regeneration. Rooted shoots were transplanted to soil and grown in a greenhouse at 18-21.5 °C and artificial light supplementation between 8:00 AM and 10:00 PM if radiation is below 150W/m^2^.

#### Copper induction of CuBe plant material

For fluorescence screening of T0 plants, 5 mm-diameter discs were placed upside-down on a 96-well plate containing 250 µl 0.1 mM CuSO_4_ per well. For spray induction, leaves were sprayed with 1 or 5 mM CuSO_4_ + 0.05% fluvius surfactant on both sides. For agar-diffusion induction, sterilized seedlings were transferred to 12-well transparent plates (VWR, #10062-894) containing MS medium (pH 5.7) and 4.5 g/L phytoagar with 0.05, 0.5, and 5 mM CuSO_4_. For watering-in-pot induction, the soil-perlite substrate was watered with 1 mM or 5 mM CuSO_4_ solution (single or 9-day application). Hydroponic induction involved transferring seedlings to 3.5 cm x 3.5 cm x 4 cm rockwool cubes pre-wetted with nutrient solution. Once the seedling roots had grown through the rockwool cubes, they were transplanted to polystyrene boards placed atop trays filled with nutrient solution. An air pump was used for oxygenation. The spacing between plants was maintained at 4.5 cm. After 15 days of growth, the nutrient solution was replaced with 1 mM CuSO_4_. Post-harvest induction used detached leaves incubated in water with 0.01-1 mM CuSO_4_ or dipped in 1 or 5 mM CuSO_4_ + 0.05% fluvius for 5 minutes followed by water incubation.

#### Total soluble protein extraction

Frozen leaves were ground in liquid nitrogen. 75 mg of powder was homogenized with ice-cold PBS buffer (80 mM Na_2_HPO_4_·7H_2_O, 20 mM NaH_2_PO_4_·H_2_O, 100 mM NaCl, pH 7.4) (1:3 w/v) and centrifuged (14,000 x g, 15 min, 4°C). The supernatant was used as a total soluble protein extract. For antibody purification, 8 g of ground tissue was homogenized in 20 mM phosphate buffer (7.4 mM NaH_2_PO_4_, 12.6 mM Na_2_HPO_4_·7H_2_O, pH 7) with 0.5 mM PMSF (Sigma-Aldrich, #78830) and 50 mM sodium ascorbate, centrifuged (10,000 xg, 15 min, 4°C), and filtered (0.22 μm) to remove debris.

#### Antibody Purification by Affinity Chromatography

The filtered protein extract was loaded onto a column with Protein A agarose resin (ABT Technology) for gravity-flow purification following the manufacturer’s protocol. Bound antibodies were eluted with 100 mM citrate buffer (pH 3). The elution fractions (250 μl each) were immediately neutralized with 37.5 μl of 1 M Tris-HCl buffer, pH 9.

#### Isolation of apoplast fluid

Leaves expressing the apoplast-tagged protein were infiltrated with ice-cold PBS under vacuum for 1 minute, followed by slow release. After removing the excess buffer, leaves were rolled, placed into a 20 ml syringe in a 50 ml tube, and centrifuged (1000 xg, 10 min, 4°C). The apoplastic fluid was concentrated 6.5-fold using 10 kDa Amicon Ultra-4 10K centrifugal filters (Millipore) after centrifugation (3,000 xg, 20 min, 4°C).

#### SDS-PAGE and Western Blot Analysis

Proteins were separated by SDS-PAGE using MES-SDS buffer (pH 7.3) on NuPAGE 10% Bis-Tris gels (Invitrogen) under reducing conditions, with equal protein loading confirmed by Bradford assay. Proteins were visualized by Coomassie blue staining. For Western blot, proteins were transferred by semi-wet blotting (XCell II™ Blot Module, Invitrogen, Life Technologies) to a Polyvinylidene Difluoride (PVDF) membrane (Amersham Hybond™-P, GE Healthcare). The membrane was blocked with 2% ECL Prime (GE Healthcare) in PBS-T (PBS with 0.1% (v/v) Tween-20). For eGFP detection, the membrane was probed with 1:10,000 anti-GFP (proteintech, 66002-1-Ig), and subsequently incubated with 1:10,000 HRP-labeled anti-mouse IgG secondary antibody (GE Healthcare). To detect the anti-SARS-CoV-2 antibody, the membrane was incubated with 1:20,000 HRP-conjugated anti-human IgG (Sigma-Aldrich, #A8792). ECL Prime Western blotting Detection Reagent (GE Healthcare) was used to detect the HRP-antibodies in a Fujifilm LAS-3000 imager for chemiluminescent detection of target proteins in the membrane.

#### Protein quantification

The total protein concentration in the soluble protein extraction or the apoplastic fluid was quantified by the Bio-Rad Protein following the manufacturer’s guidelines. For quantifying the protein of interest, samples were separated by SDS-PAGE and stained with Coomassie Blue (as described above). A BSA standard curve was generated on the same gel for quantification. Densitometry analysis of bands using ImageJ software was used to estimate the concentration of the protein of interest. Briefly, band intensities (integrated peak areas) of BSA and the target protein were measured. The BSA standard curve then allowed the estimation of the relative abundance of the protein of interest.

### ELISA

Costar EIA/RIA plates (Corning) were coated overnight at 4°C with 0.4 μg RBD (RayBiotech, #230-30162) in coating buffer (15 mM Na_2_CO_3_, 35 mM NaHCO_3_, pH 9.6) per well. After washing and blocking with 2% ECL Prime blocking agent in PBS with 0.1% (v/v) Tween-20 for 2 h at RT, plates were incubated with total protein or apoplastic fluid samples for 1 h at RT. Following washes, plates were incubated with 1:2,000 HRP-conjugated anti-human IgG (Sigma-Aldrich, #A8792) for 1 h. Plates were then washed, and o-phenylenediamine dihydrochloride SIGMAFAST™ OPD tablets (Sigma-Aldrich, #P9187) were added as the substrate for the HRP colorimetric reaction. The colorimetric reaction was stopped with 3 M HCl, and 492 nm absorbance was measured. The absorbance values plotted in the graphs are the result of subtracting the background levels of a control sample from a wildtype plant.

#### Quantitative PCR analysis

Total DNA from 100 mg leaf tissue was isolated using the DNeasy Plant Kit (Qiagen). Calibration curves for Geminino 1.0-eGFP or -n72_hγHC constructs were generated using purified plasmids mixed with 5 ng of *N. benthamiana* wildtype genomic DNA. Plasmid copy number for the curve was calculated based on the *N. benthamiana* haploid genome size (7.75 x 10^8^ bp) and DNA mass-to-length conversion (660 g/mol). DNA concentration was measured by Qubit fluorometric quantification. DNA integrity and RNA contamination were assessed by gel electrophoresis. qPCR was performed on 5 ng of total DNA (3 technical replicates) using SYBR Green dye and primers amplifying the Geminino 1.0 35S terminator-SIR region. Housekeeping gene F-BOX amplification was used for verifying equal quantities of genomic DNA in the compared samples. Ct values were interpolated against calibration curves, and the DNA copy number was normalized to the *N. benthamiana* haploid genome size.

## Supporting information

Supplementary information CuBe system

## ACKNOWLEDGEMENTS

This research was supported by the Ministerio de Ciencia e Innovación (Spain) through Agencia Estatal de Investigación (grant PID2022-141438OB-I00), the European Commission – NextGenerationEU through the Ministerio de Ciencia e Innovación (Spain) and Agencia Estatal de Investigación (grant TED2021-131171B-I00). Work in RLD’s laboratory is partially funded by the Excellence Strategy of the German Federal and State Governments. E.G.P. is recipient of ACIF-2020 fellowship (Generalitat Valenciana). The authors would like to thank Asunción Fernandez Del Carmen for her assistance with experimental design and data analysis.

## SUPPLEMENTARY INFORMATION

### Supplementary figures

**Figure S1.** Analysis of the OCS terminator (OCSt) presence in the offspring of T1 line 9.

**Figure S2.** Symptoms caused by 5 mM CuSO_4_ spraying of CuBe-eGFP leaves after four days.

**Figure S3.** CuBe-eGFP seedlings grown in MS-agar plates to monitor copper movement through the plant at time 0 post-sowing (a) and 5 days post-sowing (b).

**Figure S4.** CuBe-eGFP seedlings grown in MS-agar plates to monitor copper movement through the plant.

**Figure S5.** 5-week-old CuBe-eGFP plants grown in hydroponic conditions.

**Figure S6.** Quantification of eGFP expressed by CuBe-eGFP plants inducted by hydroponic application of 1 mM CuSO_4_.

**Figure S7.** Symptoms caused by leaf incubation from CuBe-eGFP plants line 4.11 and 7.8, and WT in CuSO_4_ solution for 2 (left) or 5 days (right).

**Figure S8.** Leaves from CuBe-eGFP plants line 4.11 and 7.8, and WT after a 5-min incubation in 1 mM CuSO_4_ + 0.05% fluvius solution and maintenance in water for 2 days.

**Figure S9.** Damage symptoms caused by incubation of leaves from CuBe-eGFP plants line 4.11 in water for 4 days after a 5-min incubation in 5 mM CuSO_4_ + 0.05% fluvius solution.

**Figure S10.** Electrophoresis gel from genomic DNA of three leaves that come from different CuBe-eGFP plants.

**Figure S11.** Leaves from 8-week-old (flowering) plants and 5-week-old plants of CuBe-eGFP line 4.11 after a 5 min pulse incubation in 1 mM CuSO_4_ + 0.05% fluvius, followed by 3 days incubation in water.

**Figure S12.** Quantification of purified n72_hγHC expressed by CuBe-n72_hγHC plants inducted by pulse 1 mM CuSO_4_ copper incubation of post-harvested leaves.

**Figure S13.** Occasional escape of replicon in isolated cells in untreated leaves from CuBe-eGFP plants.

#### Supplementary tables

**Table S1.** Genetic screening of transgenic plants regenerated with single-regulation for Rep/RepA and constitutive CUP-Gal4.

**Table S2.** Genetic screening of T0 CuBe plants.

**Table S3.** Segregation analysis of Kanamycin resistance for T1 CuBe-eGFP lines.

**Table S4.** Segregation analysis of Hygromycin resistance for T1 CuBe-eGFP lines.

**Table S5.** Segregation analysis of Kanamycin (Kan) and Hygromycin (Hyg) resistance for T1 CuBe-eGFP lines.

**Table S6.** Segregation analysis of Kanamycin resistance for T1 CuBe-n72_hγHC lines.

**Table S7.** Segregation analysis of Hygromycin resistance for T1 CuBe-n72_hγHC line.

**Table S8.** GB level 1 and >1 transcriptional units and modules. Sequences can be found at https://goldenbraidpro.com/ using the GB number.

