## Supplementary information CuBe system for "CuBe: a geminivirus-based copper-regulated expression system suitable for post-harvest activation"

**Supplementary figures**

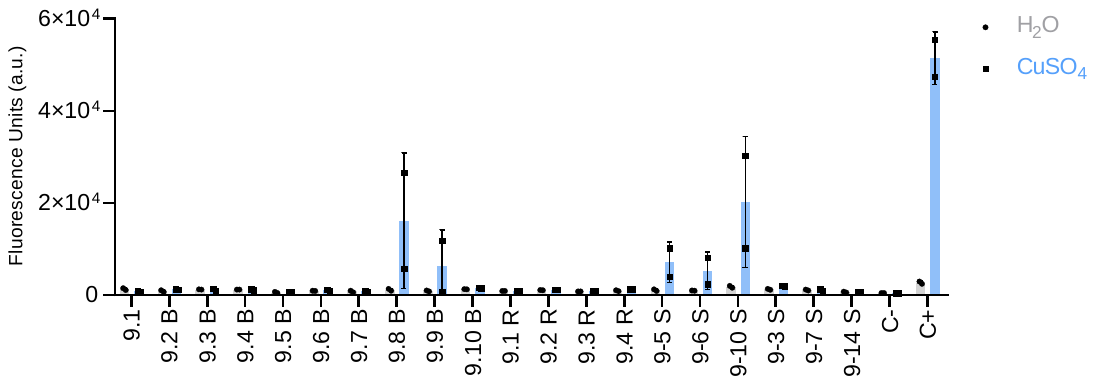

Group R

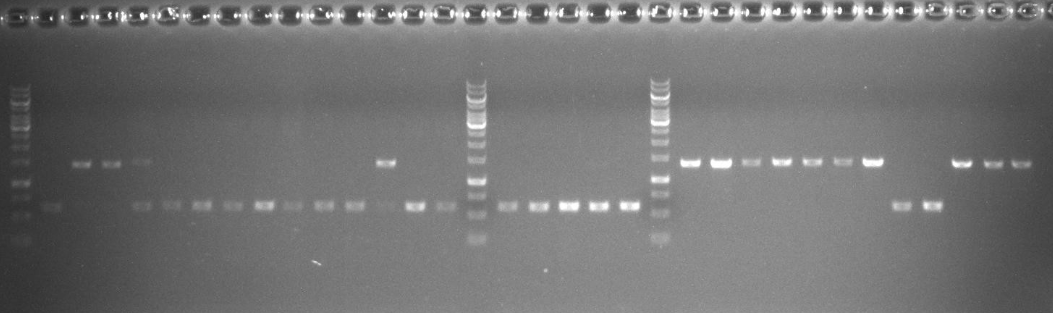

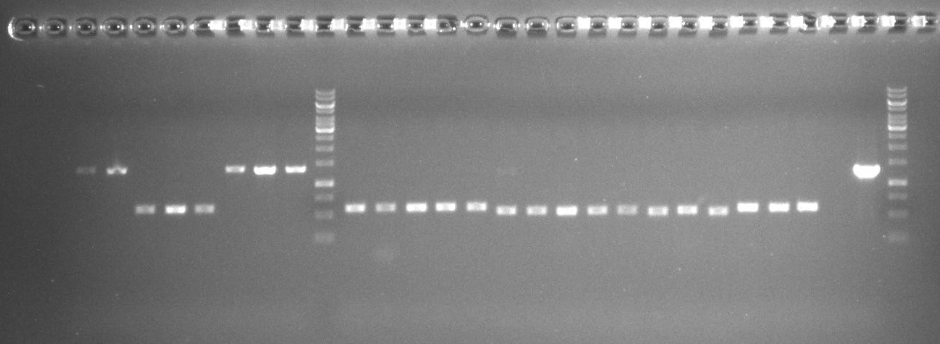

FLP, T1, line 9

Group B

1	2	3	4	5	6	7	8	9	10

Group R

1	2

3	4	5	6	10	7	3	14

Group S

FLP, T1, line 9

a

b

M

M

**Figure S1.** Analysis of the OCS terminator (OCSt) presence in the offspring of T1 line 9. **a)** Electrophoresis gel of PCR products from amplification of the region between the start codon of Rep/RepA and the distal region before the CBS operator (see Figure 3a). Only the plants where the OCSt is gone by recombination (red square) are functional, as reported in b). **b)** Expression of eGFP in T1 line 9 plants evaluated in a) after incubation of leaf discs in water or copper sulfate. C- is negative control (empty vector infiltrated in wild-type plant), and C+ is positive control (transient expression of CuBe system in wild-type plant).

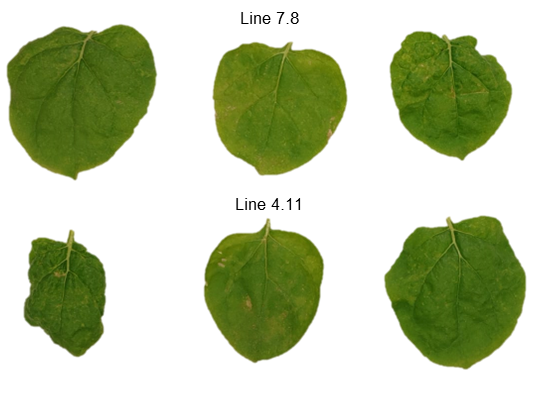

**Figure S2**. Symptoms caused by 5 mM CuSO_4_ spraying of CuBe-eGFP leaves after four days. The first row of leaves is from transgenic line 7.8, and the second row is from line 4.11.

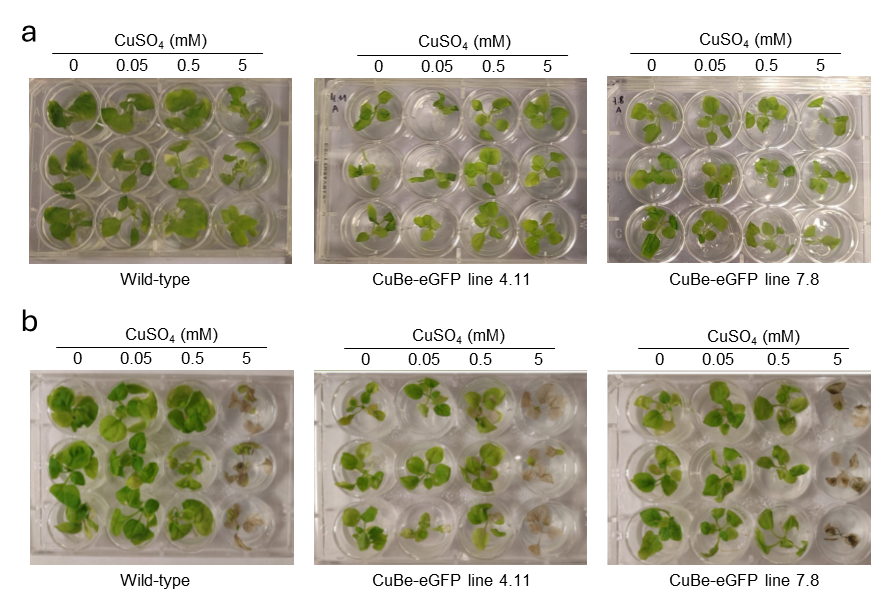

**Figure S3**. CuBe-eGFP seedlings grown in MS-agar plates to monitor copper movement through the plant at time 0 post-sowing (a) and 5 days post-sowing (b).

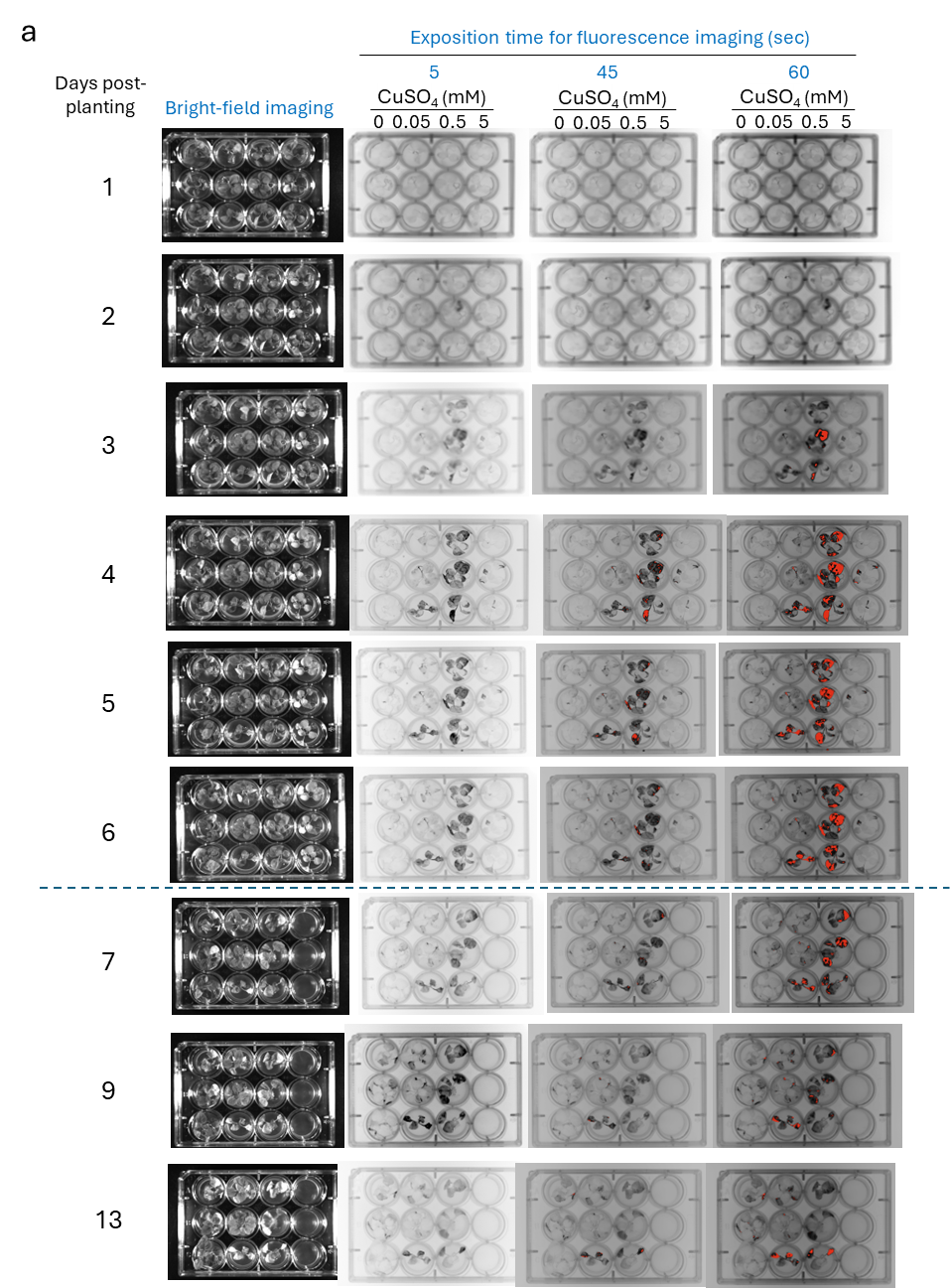

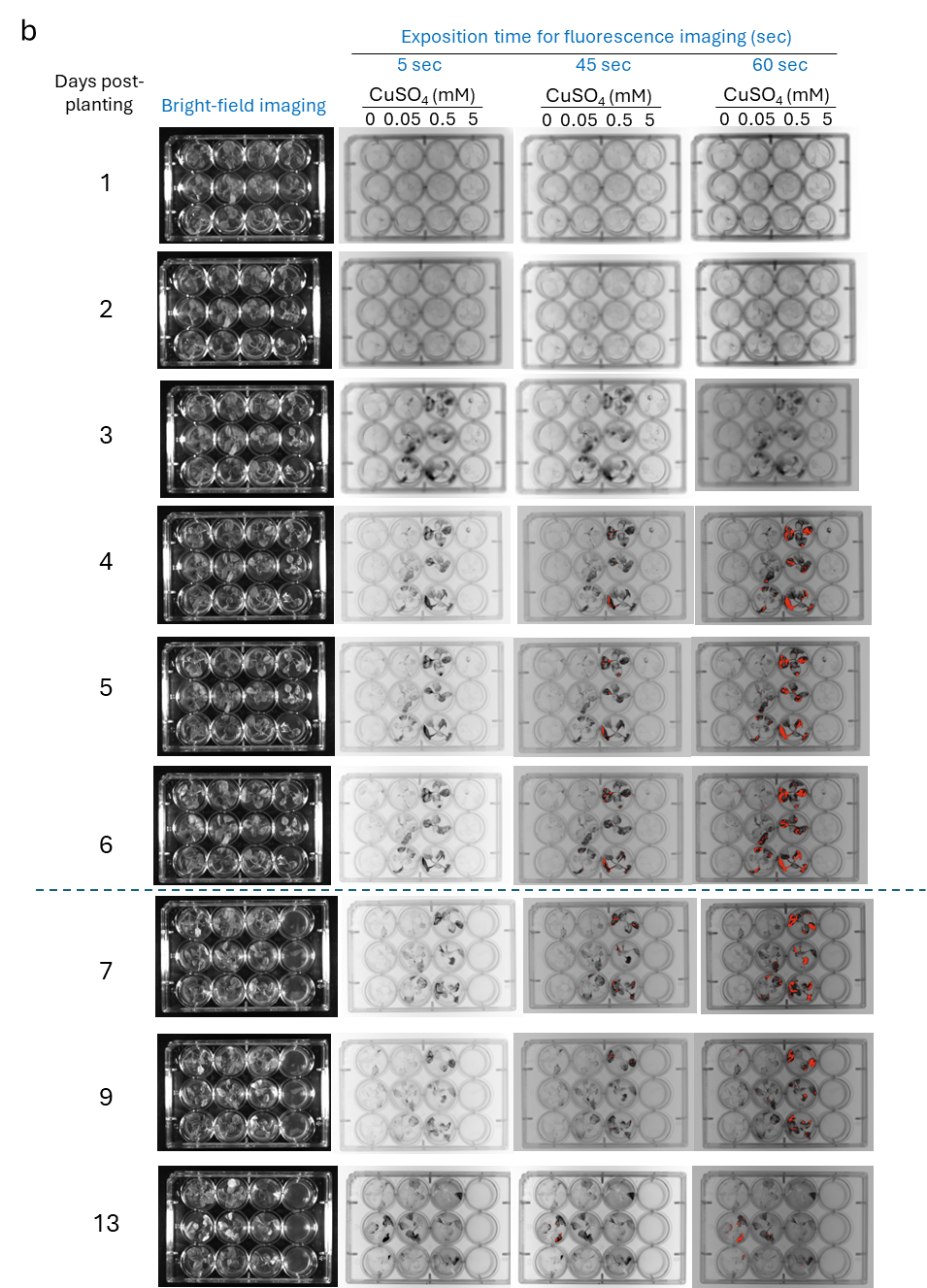

**Figure S4**. CuBe-eGFP seedlings grown in MS-agar plates to monitor copper movement through the plant. Seedlings in a) are CuBe-eGFP line 4.11, and seedlings in b) are CuBe-eGFP line 7.8. The blue line indicates a transfer of seedlings to new plates without CuSO_4_.

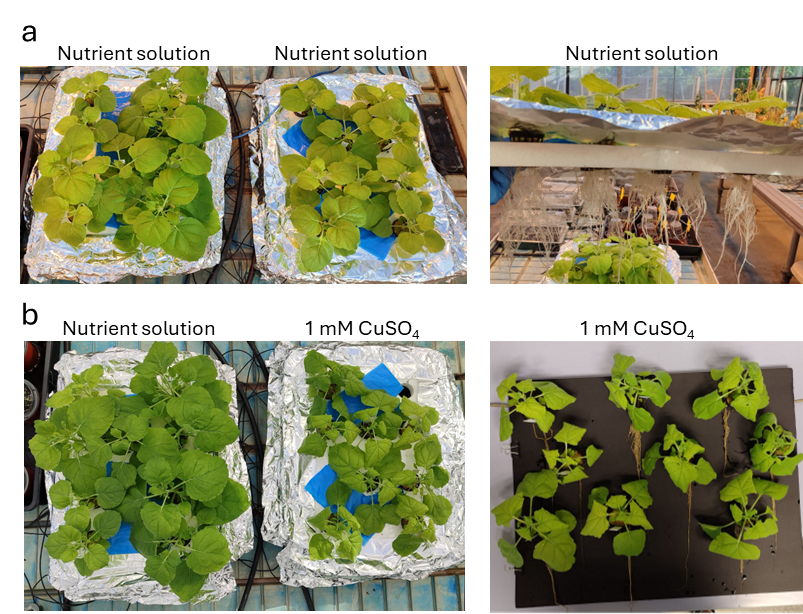

**Figure S5.** 5-week-old CuBe-eGFP plants grown in hydroponic conditions. a) Plants before exchanging the standard nutrient solution. On the right, white and healthy roots are shown. B) Plants three days after exchanging nutrient solution by 1 mM CuSO_4_ solution in the tray on the right in the picture on the left. The picture on the right shows the browning of the roots after copper treatment.

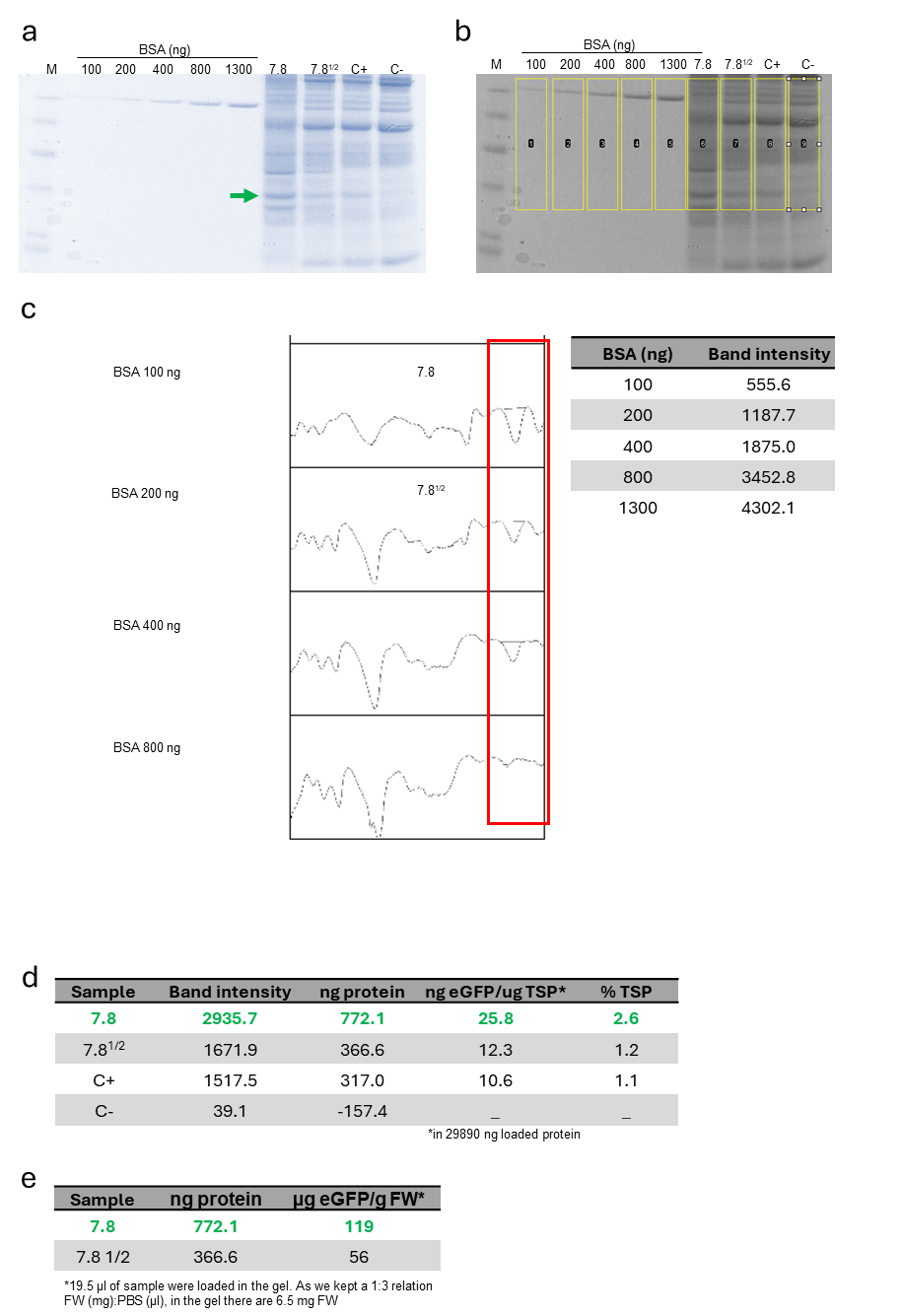

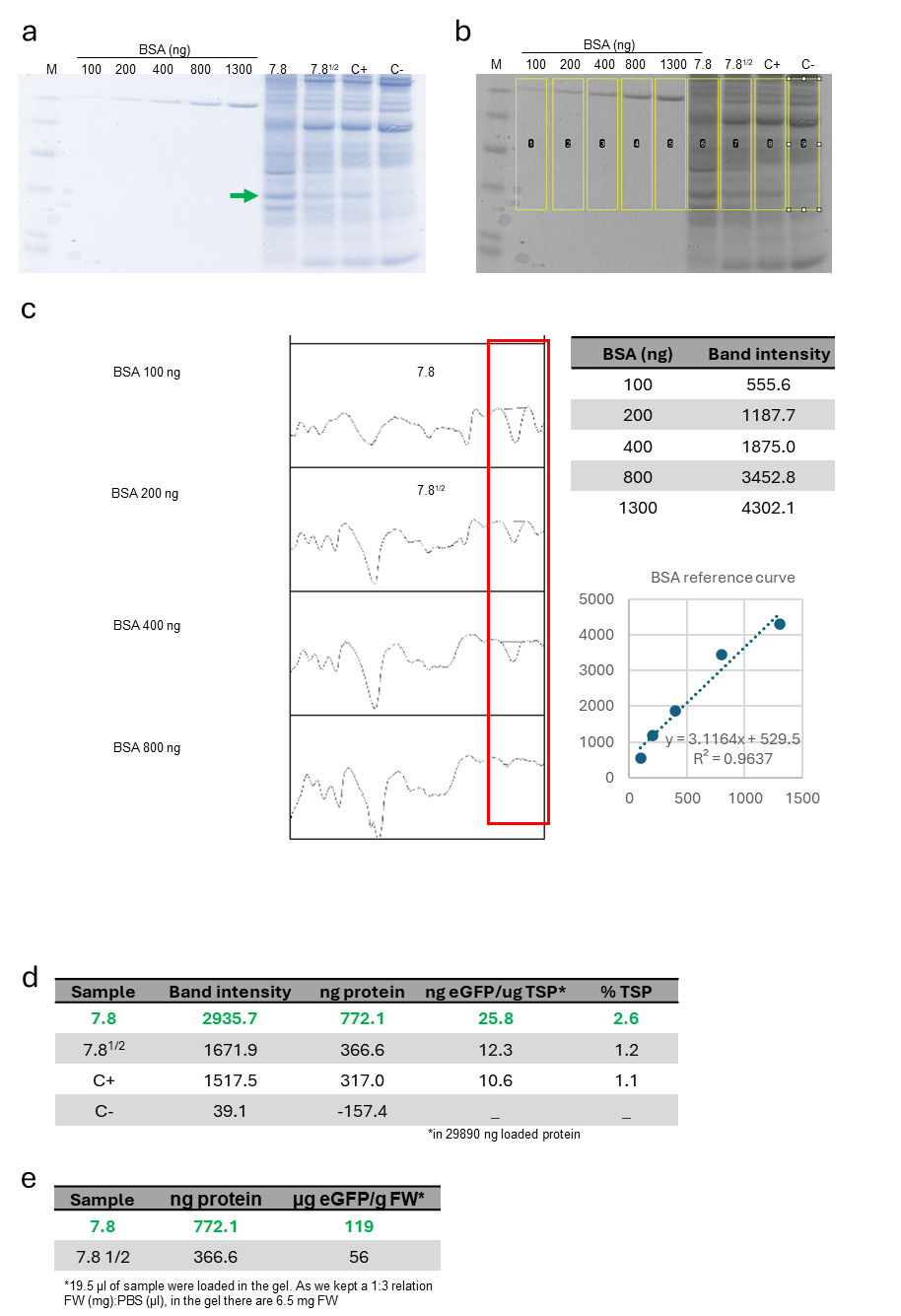

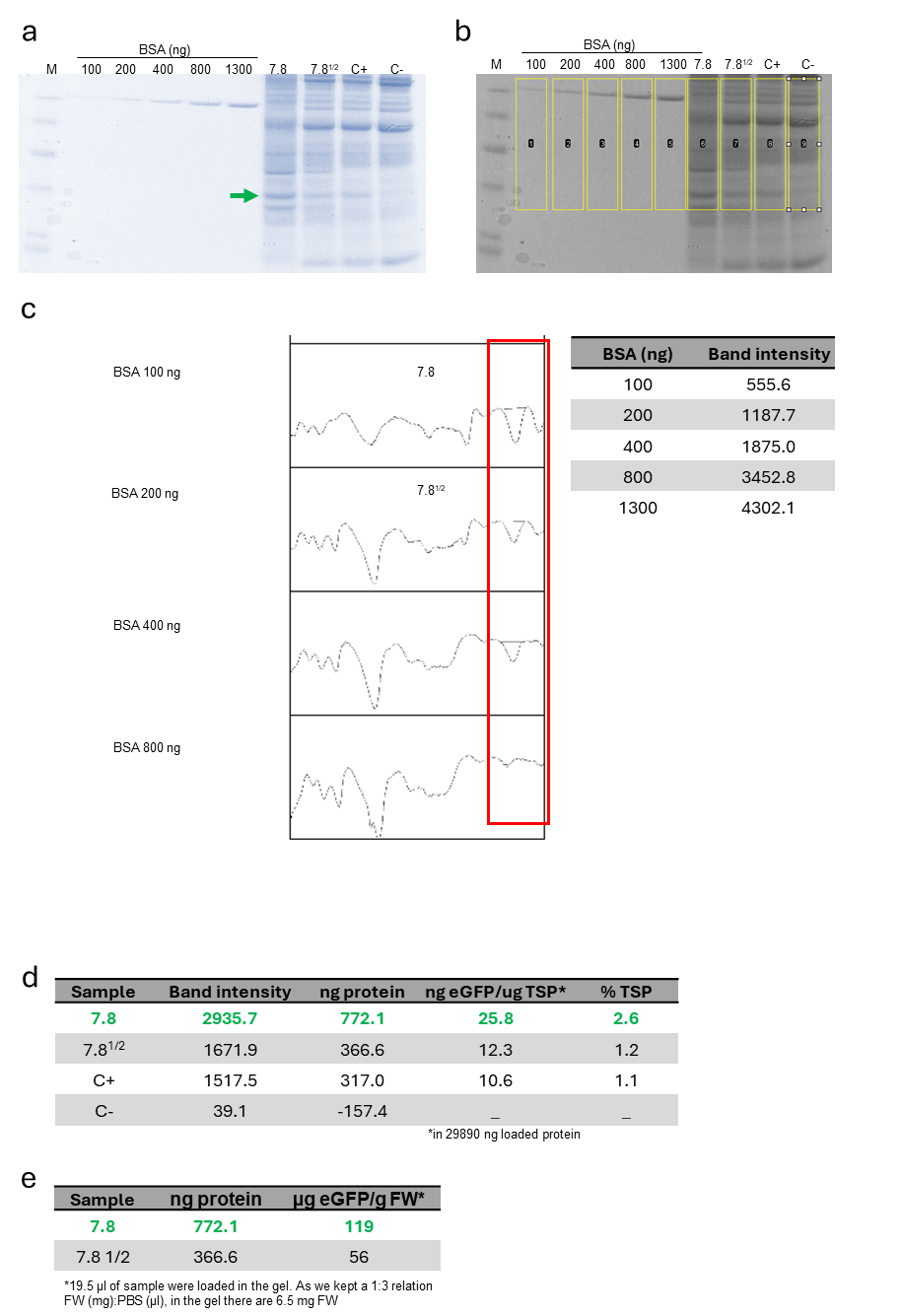

**Figure S6**. Quantification of eGFP expressed by CuBe-eGFP plants inducted by hydroponic application of 1 mM CuSO_4_. a) Coomassie-stained gel containing: BSA protein as reference for protein quantity (lanes 2-6), total soluble protein (TSP) of CuBe-eGFP line 7.8 (lane 7), same sample but diluted 1:2 with WT TSP (lane 8), positive control (C+) as TSP from transient expression of eGFP in WT plant (lane 9), and negative control (C-) as TSP from WT plant (lane 10). The green arrow points out the band corresponding to eGFP. b) Same gel as a) imaged by LAS for quantification. c) Integrated peak areas (red square) corresponding to the gel bands of BSA and the target proteins. The peak area from BSA bands and their known concentrations were used to create a reference curve to estimate the relative abundance of the protein of interest. The table on the right contains the resulting correlation between protein quantity and band intensity, which is represented in the graph below. d) Percentage of total soluble protein (% TSP) corresponding to eGFP. e) Micrograms of eGFP per gram of fresh weight of plant material (µg eGFP/g FW).

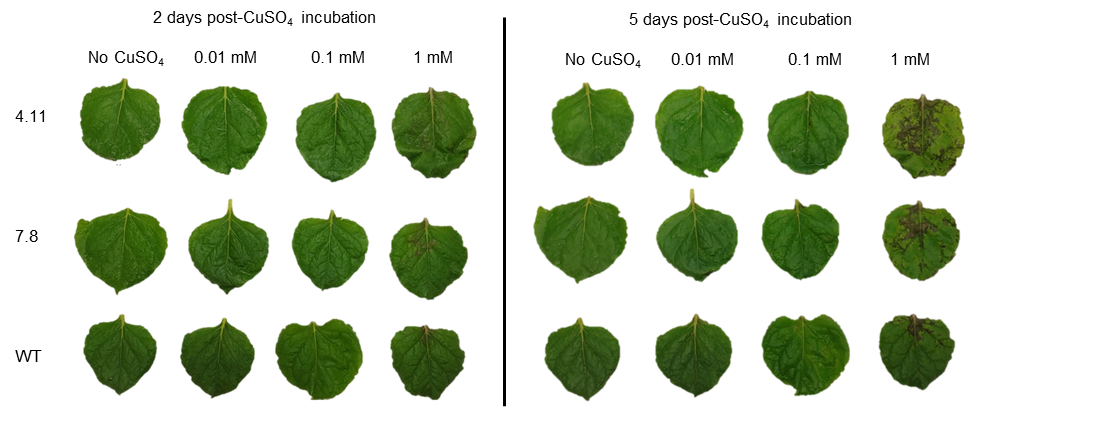

**Figure S7**. Symptoms caused by leaf incubation from CuBe-eGFP plants line 4.11 (first row) and 7.8 (second row), and WT (third row) in CuSO_4_ solution for 2 (left) or 5 days (right).

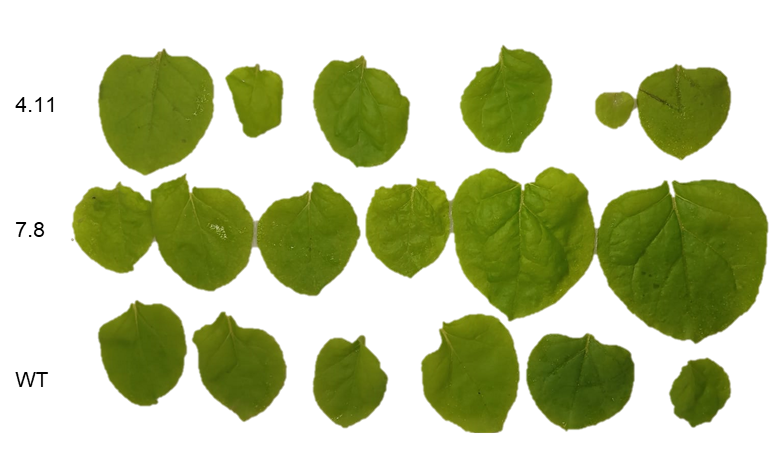

**Figure S8.** Leaves from CuBe-eGFP plants line 4.11 (first row) and 7.8 (second row), and WT (third row) after a 5-min incubation in 1 mM CuSO_4_ + 0.05% fluvius solution and maintenance in water for 2 days.

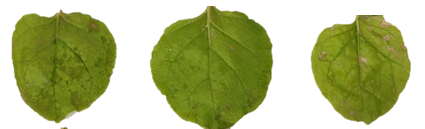

**Figure S9**. Damage symptoms caused by incubation of leaves from CuBe-eGFP plants line 4.11 in water for 4 days after a 5-min incubation in 5 mM CuSO_4_ + 0.05% fluvius solution.

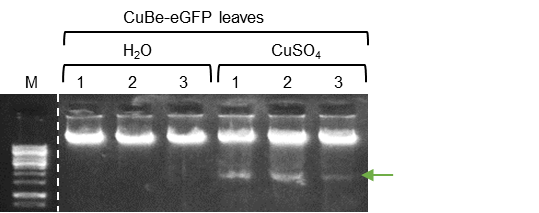

**Figure S10**. Electrophoresis gel from genomic DNA of three leaves that come from different CuBe-eGFP plants. Each number indicates one of the three assayed plants. Leaves were pulse-treated (5 min) after harvest with a CuSO_4_ + 0.05% fluvius solution and kept in H_2_0 for 3 days. The green arrow points to the band associated with the Geminino 1.0 replicative unit whose replication is inducted by CuSO_4_.

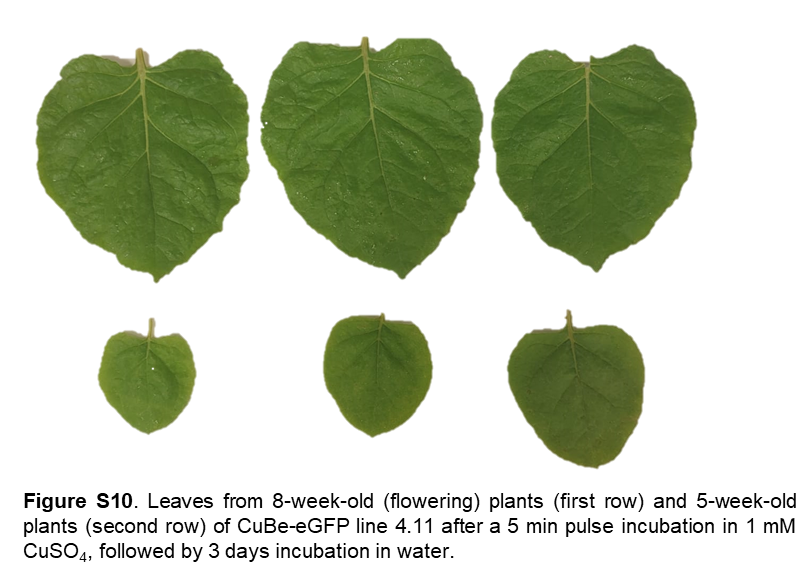

**Figure S11**. Leaves from 8-week-old (flowering) plants (first row) and 5-week-old plants (second row) of CuBe-eGFP line 4.11 after a 5 min pulse incubation in 1 mM CuSO_4_ + 0.05% fluvius, followed by 3 days incubation in water.

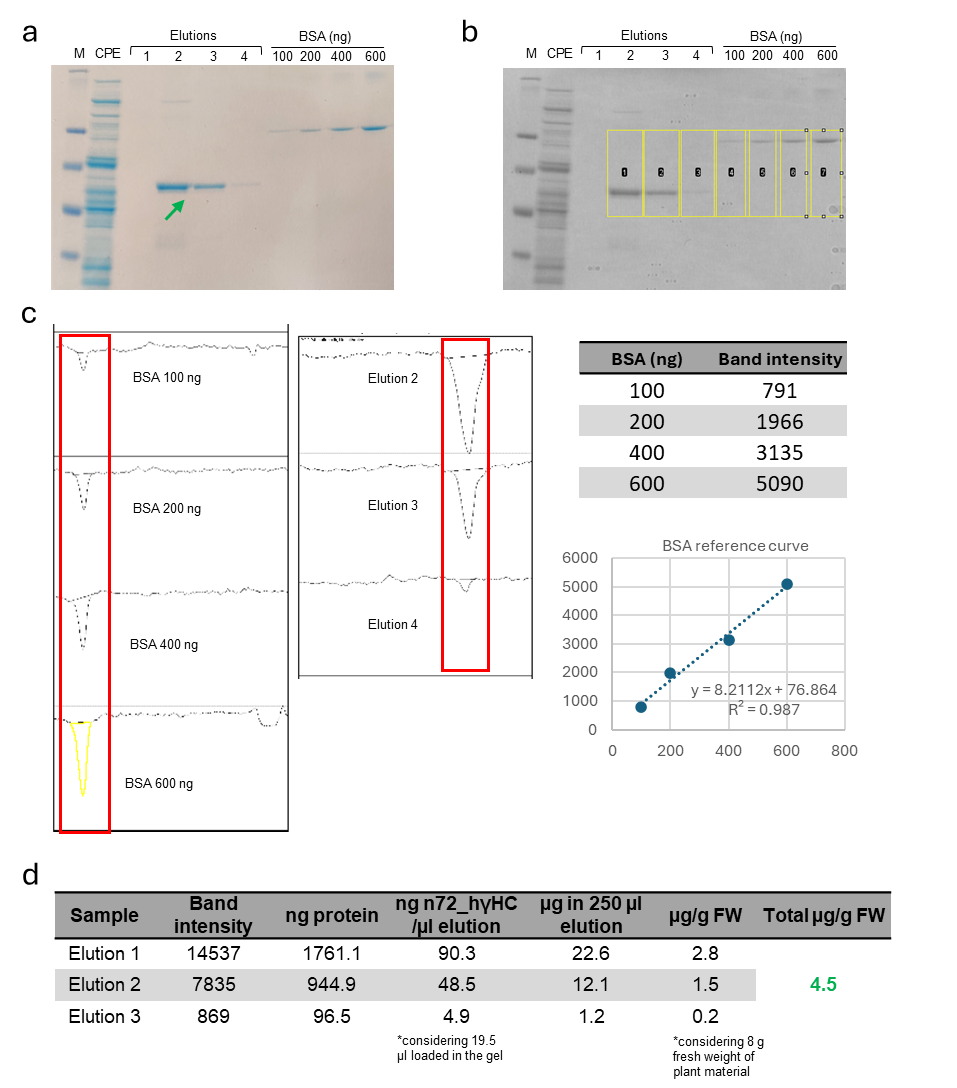

**Figure S12**. Quantification of purified n72_hγHC expressed by CuBe-n72_hγHC plants inducted by pulse 1 mM CuSO_4_ copper incubation of post-harvested leaves. a) Coomassie-stained gel containing BSA protein as reference for protein quantity (lanes 7-10), crude protein extract (CPE) of CuBe-n72_hγHC (lane 2), and elutions 1, 2, 3, and 4 from protein A purification (lanes 3-6). The green arrow points out the band corresponding to n72_hγHC. b) Same gel as a) imaged by LAS for quantification. c) Integrated peak areas (red square) corresponding to the gel bands of BSA and the target proteins. The peak area from BSA bands and their known concentrations were used to create a reference curve to estimate the relative abundance of the protein of interest. The table on the right contains the resulting correlation between protein quantity and band intensity, which is represented in the graph below. d) Micrograms of n72_hγHC per gram of fresh weight of plant material (Total µg/g FW).

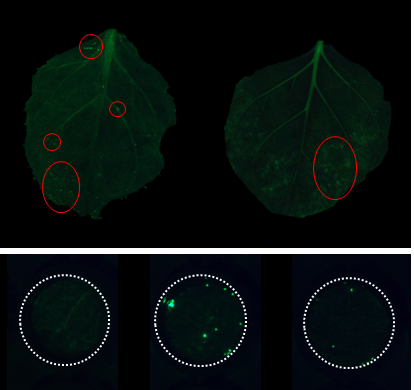

**Figure S13.** Occasional escape of replicon in isolated cells in untreated leaves from CuBe-eGFP plants. Red circles in the leaves indicate green fluorescence. The panel below shows three leaf discs where green dots might be attributed to the leaky expression of eGFP because of escaped Geminino 1.0.

**Supplementary tables**

**Table S1**. Genetic screening of transgenic plants regenerated with single-regulation for Rep/RepA and constitutive CUP-Gal4.

| **Plant line** | **CUP PCR** | **Rep PCR** | **Functionality** | **Seed production** |
| --- | --- | --- | --- | --- |
| 1 | - | + | nd | - |
| 2 | + | + | - | - |
| 3 | + | - | nd | - |
| 4 | + | + | - | - |
| 5 | + | + | - | - |
| 6 | + | + | + | - |

For the PCR columns, ‘+’ indicates positive PCR result for amplification of CUP-Gal4 or Rep/RepA as indicated by the presence of a specific band in electrophoresis analysis, and ‘-’ indicates negative PCR. ‘nd’ indicates no data. ‘+’ in functionality columns means the plant expressed eGFP upon copper induction. The negative symbol in the seed production column indicates that no plants produced viable seeds.

**Table S2**. Genetic screening of T0 CuBe plants.

| **Plant line** | **CUP** | **eGFP** | **OCSt** |
| --- | --- | --- | --- |
| 1 | + | + | + |
| 2 | + | + | + |
| 3 | + | + | na |
| 4 | + | + | - |
| 5 | + | + | -/+ |
| 6 | + | + | - |
| 7 | + | + | -/+ |
| 8a | + | + | -/+ |
| 8b | + | + | - |
| 9 | + | + | + |
| 10 | + | + | - |

In the CUP and GFP columns, ‘+’ indicates positive PCR results for CUP-Gal4 and eGFP amplification, respectively, as indicated by the presence of a specific band in electrophoresis analysis, and ‘-’ indicates negative PCR. ‘nd’ indicates no data. ‘+’ in the OCSt column indicates that the PCR product matches the expected size in electrophoresis gel if OCSt is present, ‘-’ indicates that the band matches the absence of OCSt, and ‘-/+’ indicates that the plants contain both bands matching the presence and the absence of OCSt.

**Table S3**. Segregation analysis of Kanamycin resistance for T1 CuBe-eGFP lines.

| **T1 line** | **Observed R** | **Observed S** | **Expected R (3:1)** | **Expected S (3:1)** | **P-value** | **Analysis of T-DNA insertion** |
| --- | --- | --- | --- | --- | --- | --- |
| **3** | 20 | 9 | 21.75 | 7.25 | 0.4530 | 3:1; one locus |
| **4** | 10 | 12 | 16.5 | 5.5 | 0.0014 | 1:1; one locus, insertion disrupting gene essential for male or female gamete |
| **6** | 9 | 16 | 18.75 | 6.25 | 0.0001 | 1:2; one locus, insertion disrupting gene essential for male or female gamete |
| **7** | 15 | 10 | 18.75 | 6.25 | 0.0833 | 3:1; one locus |

R indicates Resistance, S indicates susceptibility

**Table S4**. Segregation analysis of Hygromycin resistance for T1 CuBe-eGFP lines.

| **T1 line** | **Observed R** | **Observed S** | **Expected R**  **(3:1)** | **Expected S**  **(3:1)** | **P-value** | **Analysis of T-DNA insertion** |
| --- | --- | --- | --- | --- | --- | --- |
| **3** | 27 | 2 | 21.75 | 7.25 | 0.0244 | 15:1; two loci |
| **4** | 23 | 6 | 21.75 | 7.25 | 0.5919 | 3:1; one locus |
| **6** | 34 | 2 | 27 | 9 | 0.0071 | 15:1; two loci |
| **7** | 26 | 13 | 29.25 | 9.75 | 0.2294 | 3:1; one locus |

R indicates Resistance, S indicates susceptibility

**Table S5**. Segregation analysis of Kanamycin (Kan) and Hygromycin (Hyg) resistance for T1 CuBe-eGFP lines.

| **T2 line** | **Observed R-S Kan** | **Observed R-S Hyg** |
| --- | --- | --- |
| **4.11** | 38-0 | 33-0 |
| **7.8** | 25-0 | 26-0 |

R indicates resistance, S indicates susceptibility

**Table S6.** Segregation analysis of Kanamycin resistance for T1 CuBe-n72_hγHC lines.

| **T1 line** | **Observed R** | **Observed S** | **Expected R (3:1)** | **Expected S (3:1)** | **P-value** | **Analysis of T-DNA insertion** |
| --- | --- | --- | --- | --- | --- | --- |
| **6** | 24 | 10 | 25.5 | 8.5 | 0.5525 | 3:1; one locus |
| **8** | 25 | 12 | 27.75 | 9.25 | 0.2965 | 3:1; one locus |
| **14** | 38 | 6 | 33 | 11 | 0.0817 | 3:1; one locus |
| **15** | 29 | 12 | 30.75 | 10.25 | 0.5279 | 3:1; one locus |
| **16** | 33 | 7 | 30 | 10 | 0.2733 | 3:1; one locus |

*R indicates resistance, S indicates susceptibility

**Table S7**. Segregation analysis of Hygromycin resistance for T1 CuBe-n72_hγHC lines.

| **T1 line** | **Observed R** | **Observed S** | **Expected R (3:1)** | **Expected S (3:1)** | **P-value** | **Analysis of T-DNA insertion** |
| --- | --- | --- | --- | --- | --- | --- |
| **6** | 36 | 4 | 30 | 10 | 0.0244 | ~ 15:1, two loci |
| **8** | 36 | 3 | 29.95 | 9.75 | 0.0285 | ~ 15:1, two loci |
| **14** | 34 | 9 | 32.25 | 10.75 | 0.5377 | 3:1; one locus |
| **15** | 32 | 14 | 34.5 | 11.5 | 0.3946 | 3:1; one locus |
| **16** | 33 | 9 | 31.5 | 10.5 | 0.5930 | 3:1; one locus |

*R indicates resistance, S indicates susceptibility

**Table S8**. GB level 1 and >1 transcriptional units and modules. Sequences can be found at <https://goldenbraidpro.com/> using the GB number or GB name.

| **GB number** | **Construct** | **GB name** | **Description** |
| --- | --- | --- | --- |
| GB0107 | Empty vector | pEGB SF | Used as a stuffer construct to get the same infiltration DO for every *Agrobacterium* mix. |
| GB1541 | pNOS:CUP2-Gal4 | pEGB_3alpha2 PNos:Cup2:Gal4AD:Tnos | Transcriptional unit for constitutive expression of the transcriptional activator protein CUP2 in C-terminal fusion with GAL4 activation domain. Binds to specific DNA operator (CBS) in the presence of copper ions and promotes transcription. |
| GB4279 | 35S:eGFP | 3a1_35S:eGFP:T35S | Transcriptional unit for constitutive expression of the enhanced GFP. |
| GB3598 | pNOS:Rep/RepA | pDGB3_α1_pNOS:Rep/RepA:tNOS | Transcriptional unit for constitutive expression of BeYDV Rep/RepA (C1:C2). |
| GB4312 | Standard replicon | 3a1_LIR1:35S:eGFP:T35S-SIR:LIR2 | BeYDV LIR1 + TU 35S:eGFP:T35S + BeYDV SIR-LIR2 in alpha1. |
| GB5177 | INPACT replicon | o1_LIR:2nd intron:3'eGFP:T35S:SIR:p35S:5'Luc:1st intron:LIR | LIR:2nd intron + 3'eGFP +T35S.SIR + p35S + 5'Luc + 1st intron:LIR for INPACT configuration. |
| GB4480 | Geminino 1.0 | a2_NEW BeYDV geminino 3.0_eGFP | BeYDV LIR1:2nd intron + enhanced GFP CDS + Ter35S + p35S + 1st intron:LIR1 in alpha2. Deconstructed replicon that needs BeYDV Rep for circularization and, thus, replication. The intronic parts have AGGT for processing. |
| GB5178 | Geminino 2.0 | a2_geminino 4.0 (no ATG)-eGFP | Geminino 2.0-no ATG on CDS. BeYDV LIR1 + 2nd half intron ICON MP + eGFP w/o ATG + t35S:SIR + P35s + 1st half of intron from ICON MP + BeYDV LIR1. |
| GB0108 | 35S:P19 | pEGB 35s:P19:tNOS | Transcriptional unit for constitutive expression of the silencing suppressor P19. |
| GB3625 | CBS:minDFR:Rep/RepA | pDGB3_α2_4xCBS:mDFR:Rep/RepA:tNOS | Transcriptional unit for copper-regulation of BeYDV Rep/RepA with 4xCBS domain+minimal promoter DFR for induction by copper. |
| GB4646 | CBS:FRT:OCSt:FRT:minDFR:Rep/RepA | 3a1_CBS-FRT-OCS-BeYDVRepRepA-tNos | Transcriptional unit for copper- and flipase-regulated BeYDV Rep/RepA. OCSt, flanked by FRT sites, is embedded in the 5’UTR of miniDFR. |
| GB4652 | CBS:minDFR:Flp | 3a2_CBS:miniDFR:FLP:tNos | Transcriptional unit for copper-regulated flipase (Flp) recombinase. |
| GB4650 | CBS:minDFR:PhiC31 | 3a2_CBS:miniDFR:PhiC31:tNos | Transcriptional unit for copper-regulated PhiC31 recombinase. |
| GB4643 | CBS:attB:OCSt:attP:minDFR:Rep/RepA | 3a1_CBS-att-OCS-BeYDVRep/RepA-tNos | Transcriptional unit for the copper and PhiC31 regulated expression of BeYDV Rep/RepA. |
| GB4684 | Processor, amplification, pro-replicon module for eGFP | pLXB3o1 HygR-MAR10-gemiGFP-MAR10 | Module for stable transformation of Geminino 1.0-eGFP flanked by two MAR10 insulators and next to a Hygromycin resistance cassette. |
| GB4697 | Sensor module | 3a1 nptII-DISTC02-CBS-FRT:Rep/RepA:tNos-DISTC19-CBS:miniDFR:FLP:tNos-PROXC02-pNos:CUP2:Gal4AD:tNos-SF | Module for copper and flippase-regulated expression of BeYDV Rep/RepA, including the TU for copper-regulated FLP, the TU for constitutive expression of CUP2-GAL4, and the nptII cassette. |
| GB4954 | Processor, amplification, pro-replicon module for n72_hγHC | pLXB3o1_hygR:MAR10:New BeYDV geminino:SP-nano72VHH-IgG1:MAR10 | Module for stable transformation of Geminino 1.0-n72_hγHC flanked by two MAR10 insulators and next to a Hygromycin resistance cassette. |
